# Cell-state plasticity drives heterogeneity in Group 3/4 medulloblastoma

**DOI:** 10.1101/2024.02.09.579680

**Authors:** Piyush Joshi, Patricia Benites Goncalves da Silva, Konstantin Okonechnikov, Tamina Stelzer, Ioannis Sarropoulos, Mari Sepp, Jana Nolle, Mischan V. Pour-Jamnani, Anne Rademacher, Tetsuya Yamada, Céline Schneider, Julia Schmidt, Luca Bianchini, Philipp Schäfer, Kevin Leiss, Michele Bortolomeazzi, Jan-Philipp Mallm, Britta Statz, Andrea Wittmann, Kathrin Schramm, Mirjam Blattner-Johnson, Petra Fiesel, Barbara C. Jones, Till Milde, Kristian W. Pajtler, Cornelis M. van Tilburg, Olaf Witt, Karsten Rippe, Andrey Korshunov, David T.W. Jones, Volker Hovestadt, Paul A. Northcott, Marc Zuckermann, Supat Thongjuea, Natalie Jäger, Henrik Kaessmann, Stefan M. Pfister, Lena M. Kutscher

**Affiliations:** Hopp Children’s Cancer Center (KiTZ), Heidelberg, Germany; Division of Pediatric Neurooncology, German Cancer Research Center (DKFZ) and German Cancer Consortium (DKTK), Heidelberg, Germany; Developmental Origins of Pediatric Cancer Junior Research Group, German Cancer Research Center (DKFZ), Heidelberg, Germany; St. Anna Children’s Cancer Research Institute (CCRI), Austria; Cambridge Stem Cell Institute and Department of Medicine, University of Cambridge, Cambridge, United Kingdom; Center for Molecular Biology of Heidelberg University (ZMBH), DKFZ-ZMBH Alliance, Heidelberg, Germany; Centre of Genomics, Evolution and Medicine (cGEM), Institute of Genomics, University of Tartu, Tartu, Estonia; Division of Chromatin Networks, German Cancer Research Center (DKFZ) and Bioquant, Heidelberg, Germany; Single-cell Open Lab, German Cancer Research Center (DKFZ), Heidelberg, Germany; Division of Pediatric Glioma Research (B360), German Cancer Research Center (DKFZ), Heidelberg, Germany; CCU Neuropathology, German Cancer Research Center (DKFZ) and German Cancer Consortium (DKTK), Heidelberg, Germany; Department of Pediatric Oncology, Hematology & Immunology, Heidelberg University Hospital, Heidelberg, Germany; CCU Pediatric Oncology, German Cancer Research Center (DKFZ) and German Cancer Consortium (DKTK), Heidelberg, Germany; University Hospital Jena, Department of Pediatrics and Adolescent Medicine, Friedrich Schiller University Jena, Jena, Germany; Comprehensive Cancer Center Central Germany (CCCG), Jena, Germany; National Center for Tumor Diseases (NCT), NCT Heidelberg, a partnership between DKFZ and Heidelberg University Hospital, Heidelberg, Germany; Department of Neuropathology, Institute of Pathology, Heidelberg University Hospital, Heidelberg, Germany; Department of Pediatric Oncology, Dana Farber Cancer Institute, Boston, MA, USA; Broad Institute of MIT and Harvard, Cambridge, MA, USA; Division of Hematology/Oncology, Boston Children’s Hospital, Boston, MA, USA; Department of Developmental Neurobiology, St Jude Children’s Research Hospital, Memphis, TN, USA

## Abstract

Cellular heterogeneity in Group 3 and Group 4 medulloblastomas is a major driver of therapeutic intractability. Here, we describe transcription factor-driven mechanisms underlying cellular diversity in these tumors using a comprehensive single-nucleus multi-omics atlas. Our analysis reveals that rather than fixed entities, these tumors exist along a continuum of cell states defined by four molecular identity axes. We show that perturbing transcription factor activity drives cellular plasticity, a key contributor to tumor heterogeneity. Strikingly, modulation of the lineage determinant *PAX6* redirects tumor cells along both Group 3- and Group 4-like differentiation trajectories, modeling a bi-lineage medulloblastoma, and reduces aggressiveness of *MYC*-driven tumor models. Together, our findings reveal how oncogenic signals co-opt cerebellar unipolar brush cell programs to rewire tumor cell states and identify cellular plasticity as a potential targetable vulnerability in medulloblastoma.

## Introduction

Limited understanding of cellular heterogeneity in medulloblastoma, a malignant childhood cerebellar tumor group^1-3^, presents a significant therapeutic challenge. Molecular profiling over the last decade has characterized medulloblastoma into four major subgroups: WNT, SHH, Group 3 and Group 4^4^. Group 3 and 4 medulloblastomas (hereafter Group 3/4 tumors), further categorized into 8 molecular subtypes (I-VIII) spanning pure Group 3 (II, III, IV), mixed (I, V, VII), to pure Group 4 (VI, VIII) tumors^5^, together represent the most common and lethal cohort. Despite their prevalence, the inter- and intra-tumor heterogeneity of these tumors and the regulatory networks governing them remain poorly understood, impeding the development of mechanism-of-action-based therapies that could improve patient survival with reduced toxicity^6^.

Recent transcriptomic studies comparing Group 3/4 tumor gene expression programs to those of developing human cerebellum suggest these tumors arise from upper rhombic lip-derived unipolar brush cell (UBC) progenitors^7-9^. However, it remains unclear how heterogeneous Group 3/4 biology can be derived from and explained by the linear UBC differentiation process, and which regulatory networks drive malignant transformation. In this study, we generated and analyzed single-nucleus multi-omics data from 38 Group 3/4 medulloblastoma samples to provide deeper insight into the molecular mechanisms underlying the similarities and differences within Group 3/4 medulloblastoma. We focused on the differential activity of transcription-factor regulated gene regulatory networks (TF-GRNs), comprising a transcription factor and its putative downstream target genes, and identified four molecular axes of identity describing Group 3/4 medulloblastoma biology. We show that the spectrum of Group 3/4 subtypes reflects a continuum of cell states along these axes, connected through a shared regulatory landscape. We experimentally demonstrate that TF activity drives tumor cell state along these four identity axes, both *in vitro* and *in vivo*. We further show that the intermediate identity of subtype VII tumors arises from the co-existence of bi-lineage Group 3- and Group 4-like tumor trajectories originating from bi-potent precursor cells within single tumors. Finally, in a *MYC*-driven Group 3 tumor model, *PAX6* expression reduced tumor aggressiveness, a clinical phenotype also observed in *PAX6*-expressing subtype VII tumors^5^. Together, our findings provide a mechanistic framework for understanding Group 3/4 medulloblastoma biology in the context of its normal developmental origin, opening new avenues for exploring novel treatment strategies and to faithfully modeling distinct disease subtypes.

## RESULTS

### Group 3/4 medulloblastoma multi-omics atlas

When visualized in low-dimensional space, such as tSNE (t-distributed Stochastic Neighbor Embedding) or UMAP (Uniform Manifold Approximation and Projection), Group 3/4 medulloblastomas appeared as a separable yet continuous group of tumors, suggesting a gradient of biology connecting distinct molecular characteristics across tumors (bulk-RNA-seq samples, Fig. 1A; Fig. S1A-C; Table S1)^8,10-14^. Consequently, closely related tumors, such as pure Group 3 or Group 4 subtype tumors, exhibited shared enrichment of metagene programs (Fig. S1D-G; Table S2). These observations align with the previously proposed bipolar Group 3 vs Group 4 axis model of medulloblastoma biology (Fig. S1D)^10^. However, diffusion trajectory analysis of metagene programs revealed that Group 3 and Group 4 subtypes each had their own linear axis of separation (Fig. 1B; Fig. S1H-J), indicating that a multi-axial spectrum exists within Group 3/4 biology.

**Fig. 1.**
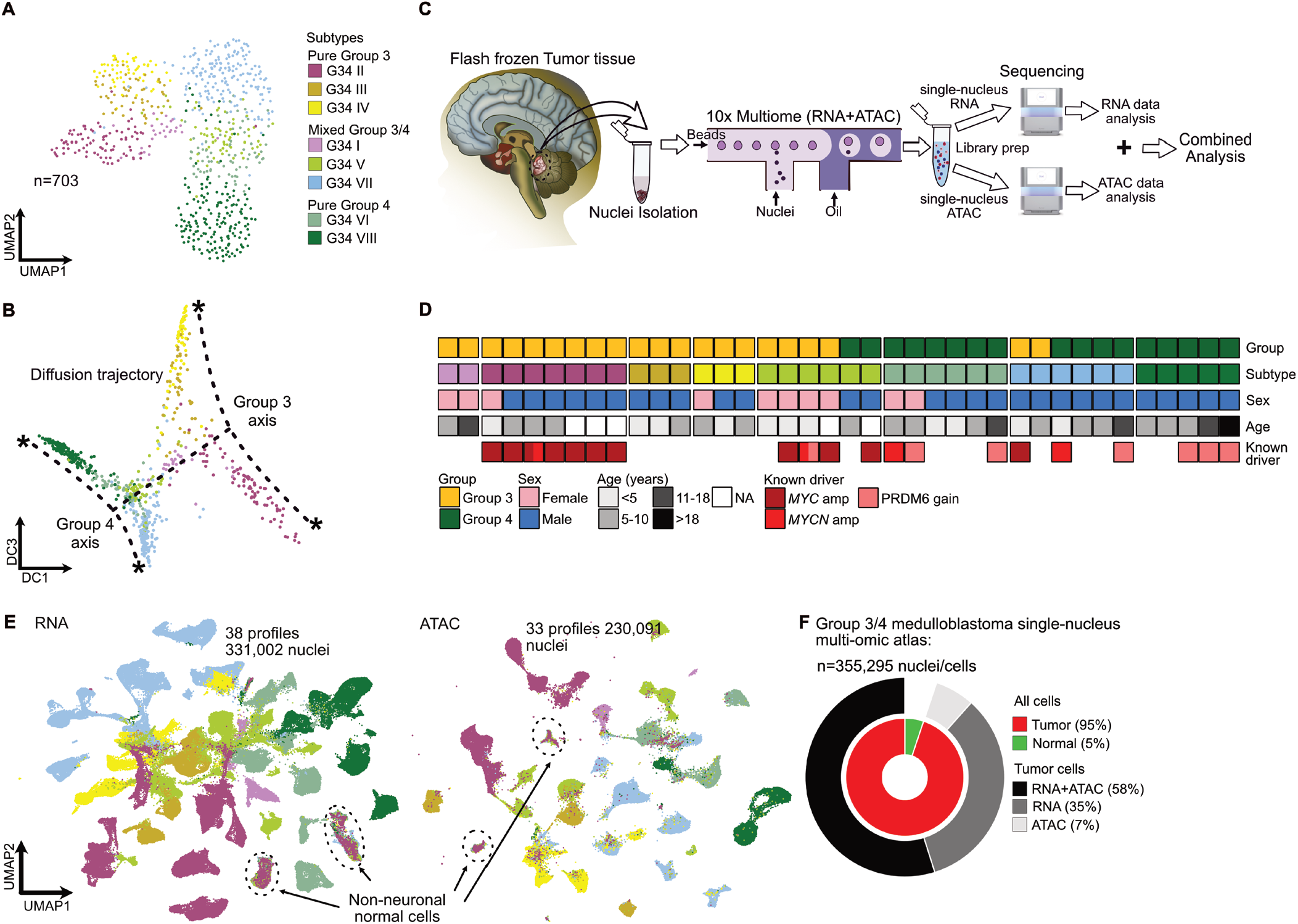
Overlapping heterogeneity defines the molecular continuity among Group 3/4 medulloblastoma. **A)** UMAP distribution of Group 3/4 medulloblastoma (n=703, bulk-RNA-seq) samples on the transcriptomic landscape colored by subtype identity. **B)** Diffusion trajectory of Group 3/4 tumors on the transcriptomic landscape. Gradients of subtype identities along the Group 3 and Group 4 axes are shown by dotted lines. **C)** Experimental design for generating single-nucleus multi-omics data from patient-derived tumor samples. **D)** Sample metadata of our Group 3/4 medulloblastoma single-nucleus multi-omics study cohort. Individual samples are represented by blocks colored per group, subtype, sex, age and known mutations. Samples with co-mutations are multicolored. **E)** UMAP distribution of snRNA-seq (left) and snATAC-seq (right) data colored by subtype identity. Non-neuronal cells are circled. **F)** Graphical summary of data modalities of single-nuclei comprising the Group 3/4 multi-omics atlas. sn-ATAC-seq data from MB248 (n=3,194 nuclei) are excluded in the chart.

We hypothesized that conserved biology across closely related subtypes is driven by shared TF-GRNs, while separable subtypes are regulated by distinct TF-GRNs. To define these molecular programs, we generated single-nucleus multi-omics data (interchangeable with “single-cell” for simplicity) from 38 Group 3/4 patient samples encompassing all eight molecular subtypes (total nuclei = 355,295; 32 samples with matched RNA and ATAC profiles, 1 sample with unmatched RNA and ATAC profiles, 5 samples with RNA only; Fig. 1C-F; Fig. S2A-H; Fig. S3A-F; Table S3). As expected, transcriptomic and chromatin accessibility profiles showed sample-specific tumor cell clusters, while normal cells, potentially representing the tumor microenvironment, integrated relatively well across samples (Fig. S2E; Fig. S3F). We used this comprehensive atlas to identify shared regulatory programs and associated cell states.

### Gene regulatory networks driving Group 3/4 identity

To integrate tumor biology within a shared regulatory framework, we transformed gene expression data into molecular program enrichment profiles. We focused on TFs with highly variable expression to identify TF-GRNs driving inter-tumor heterogeneity and continuity across Group 3/4 tumors, using a two-step approach. First, for each TF, we defined a TF-GRN within individual tumor samples using *SCENIC+* based analysis^15^, by identifying genes with expression correlated to that TF and filtering for targets with putative binding sites in associated cis-regulatory elements (CREs). We then converted the gene expression matrix into a TF-GRN score matrix and identified TF-GRNs differentially active in tumor clusters. Second, to integrate data across samples, we selected TFs associated with intra-tumor heterogeneity across multiple samples and derived a shared TF-GRN for each based on recurrent TF-target associations. TF-GRN scores were then computed for each TF across tumor cells in the integrated data and used these scores for downstream analyses, including generation of an integrated TF-GRN enrichment map (Fig. S4A; see *Methods* for details).

Using gene-set activity scores for 108 TF-GRNs (Table S4) selected from those active across tumor samples, we integrated tumor cells based on shared biology (Fig. 2A,B). The integrated tumor cell atlas revealed four axes in diffusion map space, which we labeled tumor photoreceptor-like (PR_t_), MYC-enriched, Precursor-like and tumor UBC-like (UBC_t_), based on the known function of associated TFs and enrichment of molecular programs in annotated cells, as described in detail below. Group 3 and Group 4 tumor cells contributed differently to these four axes (Fig. 2B).

**Fig. 2.**
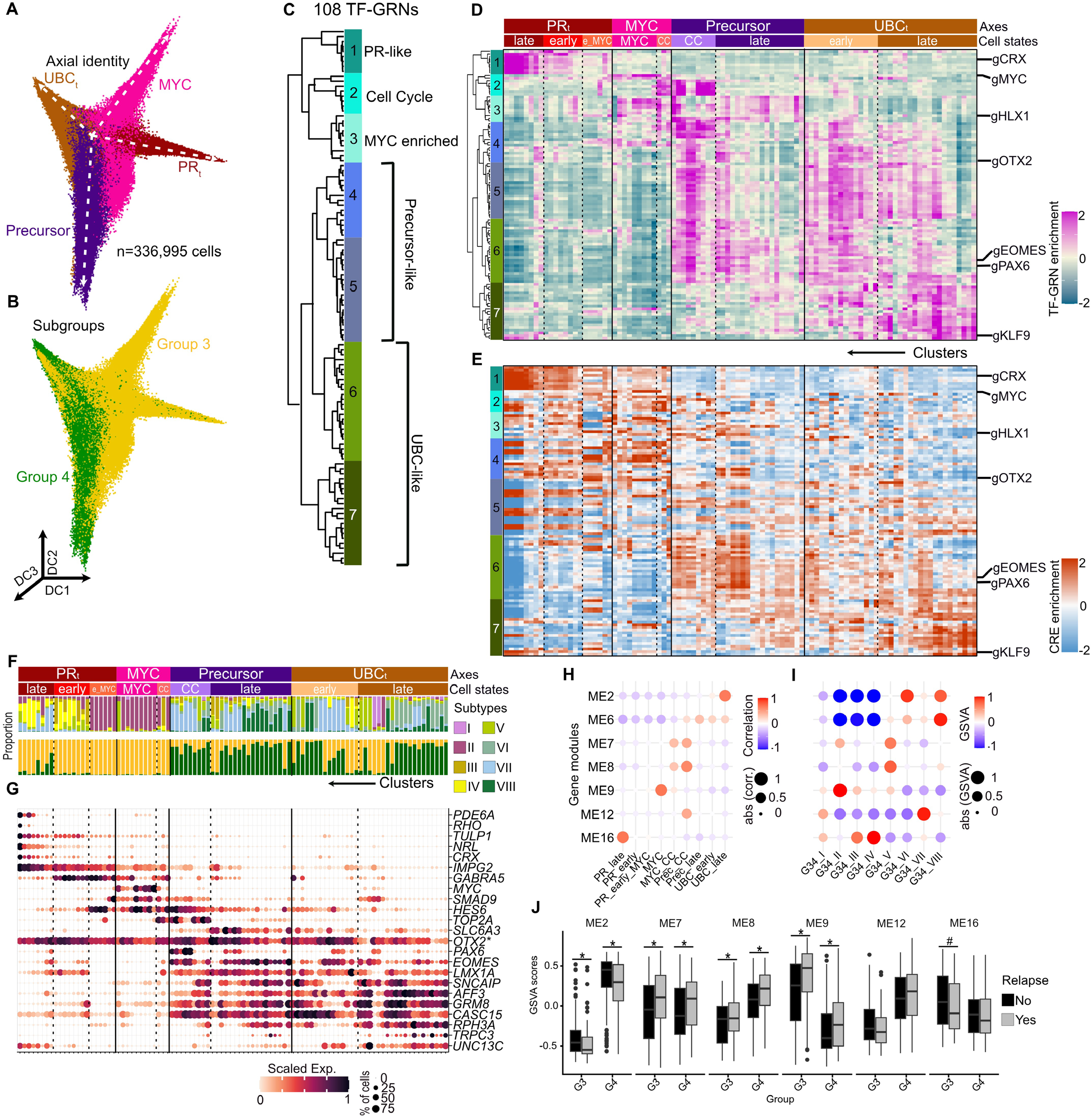
Four axes of Group 3/4 medulloblastoma identity. **A**) 3D diffusion map of Group 3/4 tumor cells obtained from TF-GRN enrichment colored by axial identity. Dotted line indicates axial trajectories. **B)** 3D diffusion map of Group 3/4 tumor cells colored by group identity. **C)** Hierarchical clustering of 108 TF-GRNs based on co-enrichment in tumor clusters. **D)** Differential enrichment of TF-GRN score across tumor clusters in the integrated data. **E)** Differential enrichment of activity of constituent cis-regulatory elements (CREs) of the TF-GRN-sets across cell clusters in the integrated data. **F)** Subtype and subgroup identity of cells comprising the cell cluster in the integrated atlas. Each bar represents a cluster’s proportional tumor subtype (top) or subgroup (bottom) composition. **G)** Marker gene expression distribution in the integrated atlas. Photoreceptor, progenitor or UBC cell-states marker genes are annotated as such. *OTX2* (marked with asterisk) is a marker gene for both photoreceptor and UBC lineages. Dot size indicates the proportion of cells in a cluster expressing a gene, and color denotes mean expression scaled across clusters per gene. **H**) Module-trait correlation between WGCNA identified modules eigengenes (y-axis) and Group 3/4 tumor cell-states (x-axis). Color and size of dot represents correlation score. Selected modules are shown. Full list of modules in Fig S4B,C,E and Table S6. **I**) Mean GSVA enrichment score of selected module eigengenes (y-axis) in tumors grouped by subtype identify (x-axis). Color and size of dot represent GSVA score. **J**) Boxplot distribution of GSVA scores of selected modules in primary and relapsed tumors grouped by subgroup identity. Data derived from https://r2.amc.nl/ for Medulloblastoma primary-relapse cohort, Korshunov, n=643, group 3/4 n=435. Asterisk represent p-value <0.05 in one-sided Wilcoxon test. # Represent p-value =0.057 in one-sided Wilcoxon test.

To molecularly define these axes, we clustered tumor cells and identified enriched TF-GRNs in each cluster. The 108 TF-GRNs were grouped into seven co-enrichment groups (Fig. 2C; Table S5). TF-GRN program 1 included well-known regulators of the photoreceptor lineage^16^ (CRX, NRL; Table S5). Programs 2 and 3 were enriched for cell-cycle and progenitor-associated TFs (MYC, E2F7, have intermediate Group 3/4 identity, were distributed across all four axes. Expression of marker genes of retinal photoreceptor (e.g., *CRX, NRL*), cycling progenitor (e.g., *TOP2A*) and cerebellar UBC (e.g., *EOMES, LMX1A*) lineages further validated these axial and cell-state annotations (Fig. 2G).

We next identified coordinated gene expression modules underlying Group 3/4 heterogeneity (Fig S4B-D, Table S6). These modules showed axial and cell-state enrichment with distinct biological associations: ME2 JUND; Table S5)^17,18^. Program 6 and 7 included well-(UBC_t_ late) was linked to NTRK signaling and synapse known regulators of early UBC development (OTX2, EOMES, LMX1A, PAX6, TWIST1; Table S5)^19,20^. We then grouped tumor clusters with similar program enrichment along each axis and subdivided them into cell states based on co-enrichment of programs defining more than one axis (Fig. 2D). Differential enrichment of CREs associated with these TF-GRNs showed largely concordant patterns (Fig. 2E), with one notable exception: along the PR_t_ axis, progenitor-like programs (2 and 3) were downregulated while their associated CREs remained comparatively accessible to those in undifferentiated MYC-axis clusters (Fig. 2D,E).

Tumor subgroup and subtype identity were also distinctively associated with the four axes. The PR_t_ and MYC axes were almost exclusively populated by Group 3 tumor cells, while the Precursor and UBC_t_ axes were predominantly populated by Group 4 tumor cells (Fig. 2F). At the subtype level, subtypes III and IV were enriched along the PR_t_ axis, while subtype II was enriched along the MYC axis (Fig. 2F). Group 4 subtypes VI and VIII showed near-exclusive association with Precursor and UBC_t_ states. Subtypes I, V and VII, which formation; ME7 and ME8 (MYC cell cycle (CC) and Precursor CC) were associated with cell cycle and DNA repair, respectively, ME9 (MYC) with ATP biogenesis; ME16 (PR_t_ late) with visual perception; and ME12 (Precursor CC), while lacking specific term enrichment, included *PAX6* in its gene-set (Fig. 2H, Table S6). Module enrichment across subtypes correlated with their cell-state composition (Fig 2F,I; Fig S4E). Comparing primary and relapsed tumors, proliferation-associated modules (ME7/8) and the MYC module (ME9) were enriched in relapsed tumors, while differentiation-associated modules (ME2/16), particularly UBC differentiation module (ME2), were enriched in primary tumors (Fig 2J; Fig S4F). This pattern was consistent in comparisons of primary-relapse pairs (Fig. S4G), suggesting that relapsed tumors, whether through tumor evolution or treatment effects, shift towards a more proliferative, less differentiated state.

In summary, the differential TF-GRN activity defines the continuum of Group 3/4 medulloblastoma biology across eight subtypes along the four axes of molecular identity.

### Mutually repressive TF-GRN interactions drive Group 3 versus Group 4 separation

To investigate transitions between the four axial identities, we examined correlations between TF-GRN activities. We hypothesized that co-expressed TF-GRNs would show high positive correlation, while mutually exclusive, potentially repressive, TF-GRN interactions would be negatively correlated (Fig. S5A). Broadly, PR_t_, MYC- and UBC_t_-associated TF-GRNs were negatively correlated, while Precursor and UBC_t_ TF-GRNs were positively correlated (Fig. 3A). In particular, gCRX/gNRL (PR_t_ axis) and gEOMES/gLMX1A (Precursor/UBC_t_ axis) were strongly negatively correlated (Fig. 3A, inset). These anti-correlative relationships suggest mutual exclusivity between PR_t_, MYC and UBC_t_ axial identities: individual tumor cells cannot simultaneously adopt two or more of these identities.

**Fig. 3.**
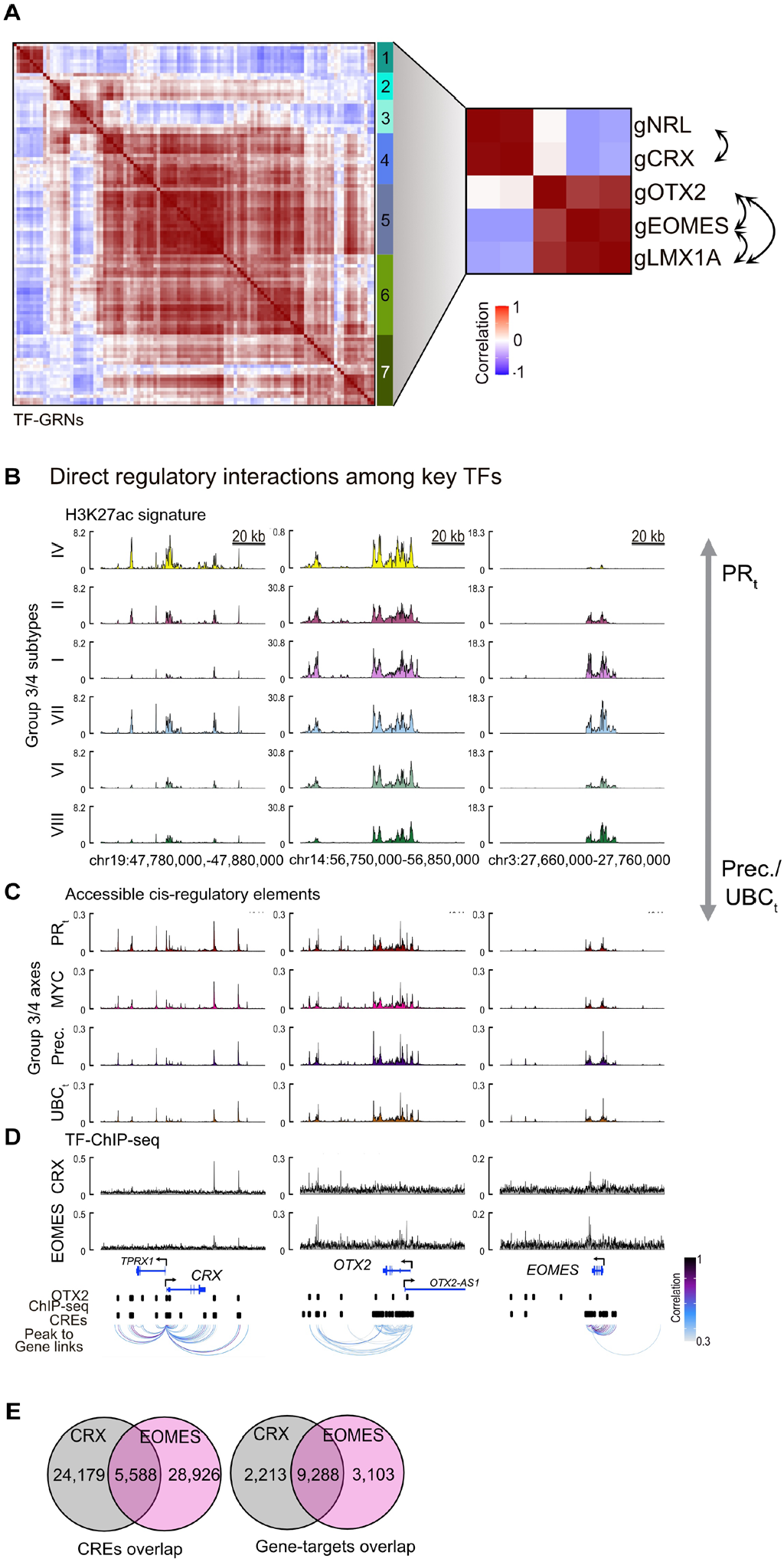
Mutually repressive PR_t_ and UBC_t_ associated TF-GRNs drive Group 3 and Group 4 identity apart. **A**) Pearson correlation analysis of TF-GRN scores in the integrated single-cell tumor data. PR_t_, MYC and UBC_t_ associated TF-GRNs show high anti-correlation. Inset, correlation between key TF-GRNs: gNRL, gCRX, gOTX2, gEOMES and gLMX1A. Arrow, putative direct interaction between TF pairs based on *SCENIC*+ analysis. Arrowhead, target of the interaction. **B)** H3K27ac ChIP-Seq^11^ shows distinct enhancer signature enrichment at *CRX, OTX2*, and *EOMES* loci across Group 3/4 subtypes. Subtypes are arranged from pure high PR_t_ (top) to high Precursor (Prec.)/ UBC_t_ (bottom) phenotype. **C)** Integrated snATAC-seq reveals differential accessibility of CREs at *CRX, OTX2*, and *EOMES* loci across Group 3/4 medulloblastoma axial identities (this study). **D)** Average ChIP-Seq (n=3) signal for CRX and EOMES (this study) and peaks for OTX2^21^ overlap CREs positively associated with expression of key genes: *CRX* (left), *OTX2* (middle) and *EOMES* (right). Interaction arcs depict representative peak to gene links colored by correlation of peak accessibility and gene expression. **E)** Venn diagram depicting overlap between summit regions (left) or associated putative target genes (right) identified from CRX and EOMES ChIP-Seq.

Next, we performed network analysis of the TF interaction graph to identify TF communities and key nodal regulators (Table S5). Globally, OTX2 and POU2F1 represented important nodes based on the number of connections to other TFs. Within TF communities, we identified local hubs based on the number of regulatory connections to other TFs within the same community (Fig S5B; Table S5). CRX and EOMES/LMX1A emerged as important regulatory hubs within the PR_t_ and Precursor/UBC_t_ TF-communities, respectively.

Taken together, these analyses suggest that TF communities, with CRX and EOMES/LMX1A playing nodal roles, are key drivers of PR_t_ and UBC_t_ separation, likely through mutual repression of their associated TF-GRN programs. To test direct regulatory interactions between CRX/NRL and EOMES/LMX1A, we examined enhancer regions around the *CRX, NRL, OTX2, EOMES*, and *LMX1A* loci using our snATAC-seq atlas; we also generated chromatin immunoprecipitation sequencing (ChIP-seq) data to map direct binding sites of CRX and EOMES in primary patient samples. By overlaying active enhancers (H3K27ac signal) in Group 3/4 tumors^11^, CRE accessibility (this study), TF binding sites for CRX and EOMES in Group 3/4 tumors (this study), and OTX2 binding in human retina^21^, we identified putatively functional CREs mediating cross-talk among these key TFs (Fig. 3B-D; Fig. S5C-E).

We found that CRX and EOMES bind to each other’s functional regulatory regions within the same tumor sample (Fig 3B-D). Combined with their anti-correlated activity (Fig 3A), this data supports a mutual inhibitory feedback loop between these factors. CRX and EOMES binding sites were enriched for their respective known binding motifs (Fig. S5F). Notably, CRX motifs, or potentially those of OTX2, which shares binding motifs with CRX, were co-enriched with EOMES in EOMES-bound regions, but not vice versa. Although some overlap was observed between CRX- and EOMES-bound regions, their putative target genes showed considerably more pronounced overlap (Fig. 3E), suggesting that the feedback between EOMES and CRX extends beyond mutual inhibition to the level of shared downstream target genes.

Given its high expression across all tumor cell states (Fig. 2G), its positive correlation with both CRX and EOMES TF-GRNS (Fig. 3A), its nodal role in integrated TF-network (Table S5), and its established roles in promoting photoreceptor identity^22^, and medulloblastoma progression^23,24^, we propose that OTX2 acts upstream of CRX/NRL to create a permissive environment for tumor cells to differentiate along both the PR_t_ and UBC_t_ lineage, while the mutually repressive interaction between CRX/ NRL and EOMES/LMX1A drives the trajectories apart.

### Cellular plasticity drives heterogeneity in Group 3/4 tumors

The distribution of tumor cell states across subtypes and individual tumors, together with the differential TF activity enrichment in these states (Fig. 2D,F), suggests that inter- and intra-tumor heterogeneity may arise from TF-driven cellular plasticity. Moreover, because most TFs are not restricted to a single cell state, their regulatory activity is likely context-dependent. To functionally validate these hypotheses, we modulated expression of selected TF genes: *MYC, CRX* and *EOMES* in two *MYC*-driven subtype II tumor models, HDMB03^25^ and MB3W1^26^, which differ in their baseline gene expression programs (Fig. S6A). For example, Precursor/UBC_t_ associated genes such as *EOMES* and *LHX1* are highly expressed in MB3W1 but are barely detectable in HDMB03 (Fig. S6A), suggesting these models may be primed to respond differently to similar perturbations.

First, we examined the effect of *MYC* knockdown in HDMB03 *in vitro*, as confirmed by Western blot (Fig. 4A), changes in morphology (Fig. S6B) and reduced proliferation (Fig. S6C). Viral manipulation alone had no effect on gene expression relative to the unmodified parental line (Fig. S6D). *MYC* knockdown resulted in upregulation of the photoreceptor-specific TF CRX, the photoreceptor-specific nuclear receptor protein NR2E3, and PR_t_ signature gene-sets (Fig 4A,B), suggesting differentiation towards the photoreceptor lineage. Knockout of *CRX*, which is normally expressed at low levels in HDMB03, did not elicit significant gene expression changes (Fig 4A; Fig S6E,F). In contrast, *EOMES* overexpression, with or without *MYC* knockdown, upregulated Group 4-specific genes while downregulating PR_t_ signature (Fig 4A,C,D). Following xenotransplantation, *EOMES* overexpression strongly increased the UBC_t_ signature with a modest gain in PR_t_ signature in resulting tumors (Fig. S6G). Together, these results indicate that both *MYC* downregulation and *EOMES* overexpression perturb the *MYC*-driven state, with *EOMES* biasing cells toward a Group 4-like identity.

**Fig. 4.**
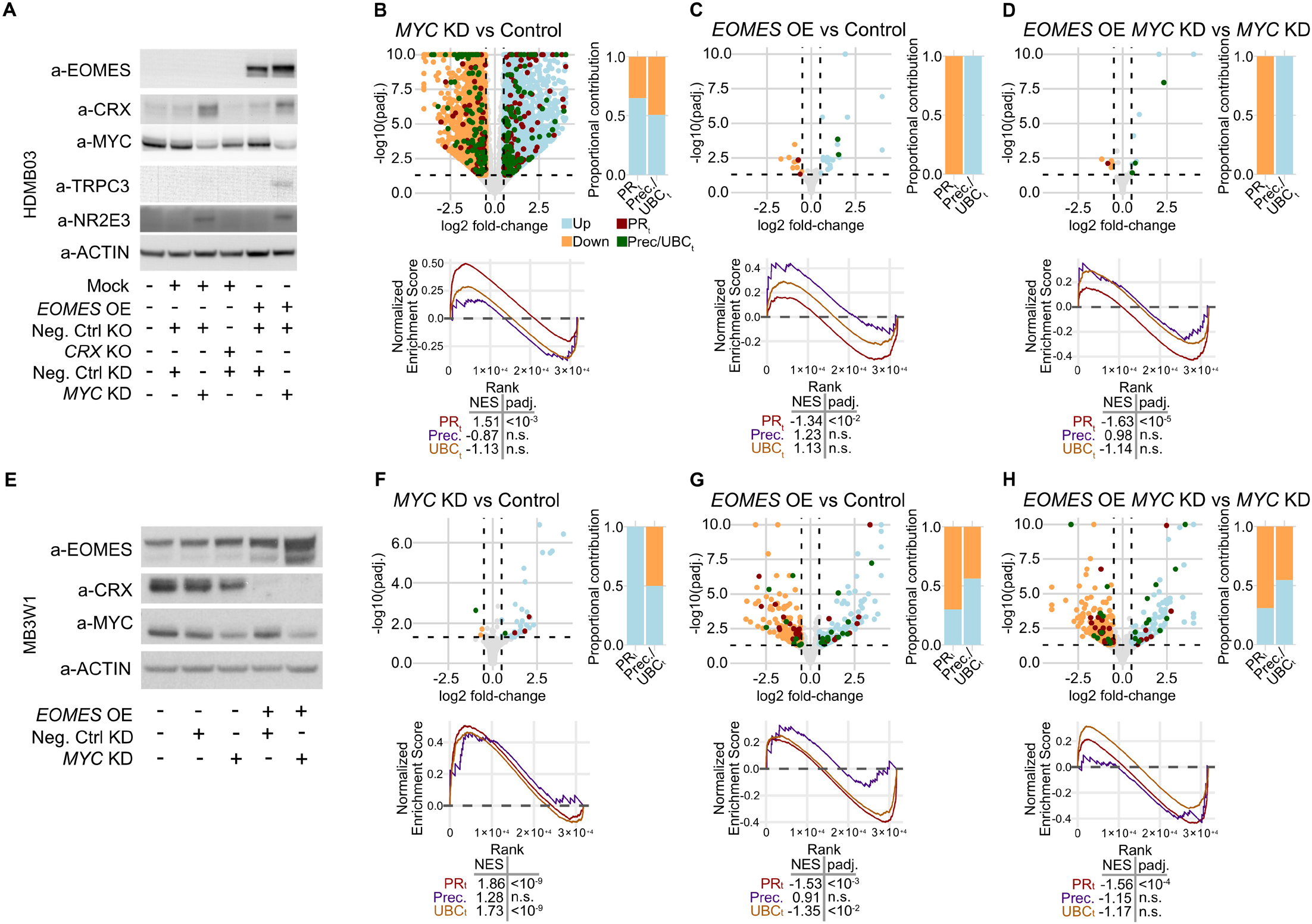
TF-driven cell states change in Group 3/4 tumors. **A)** Western blot of key proteins after modulation of *MYC, EOMES*, and *CRX* in HDMB03 *in vitro*. ACTIN, loading control. OE, overexpression. KO, knock out. KD, knockdown. **B-D)** Gene expression changes in response to TF-expression modulations in HDMB03 *in vitro*. Left: Volcano plot depicting significantly upregulated (blue) or downregulated (orange) genes in *MYC* knockdown (KD) versus control (B), *EOMES* overexpression (OE) versus control (C) and *EOMES* OE with *MYC* KD versus *MYC* KD (D). Green dots, Group 4 gene-set (Precursor (ME12) and UBC_t_ (ME2) signature genes). Red dots, Group 3 gene-set (PR_t_ (ME16) signature genes). Left: Bar plot depicting proportion of up- or down-regulated Group 3 and Group 4 signature genes that pass the significance thresholds (green or red dots in volcano plot). Bottom: Gene-set enrichment analysis (GSEA) of PR_t_, Precursor and UBC_t_ gene-set in each comparison. Table shows Normalized Enrichment Signature (NES) and padj. values for each gene-sets. **E)** Western blot of key proteins after modulation of *MYC* and *EOMES* in MB3W1 *in vitro*. ACTIN, loading control. OE, overexpression. KO, knock out. KD, knockdown. **F-H)** Gene expression changes in response to TF-expression modulations in MB3W1 *in vitro*. Left: Volcano plot depicting significantly upregulated (blue) or downregulated (orange) genes in *MYC* knockdown (KD) versus control (F), *EOMES* overexpression (OE) versus control (G) and *EOMES* OE with *MYC* KD versus *MYC* KD (H).

In MB3W1, *MYC* knockdown upregulated both PR_t_ and UBC_t_ signature gene sets, both *in vitro* and *in vivo* after xenotransplantation (Fig 4E,F; Fig S6H), in contrast to the PR_t_ dominant response observed in HDMB03. *In vitro, EOMES* overexpression downregulated both PR_t_ and UBC_t_ signature (Fig. 4G), whereas in transplanted tumors, *EOMES* overexpression drove upregulation of both (Fig S6I). In the context of *MYC* knockdown, *EOMES* overexpression attenuated the *MYC* knockdown-driven induction of the PR_t_ program *in vitro* (Fig. 4H).

Together, these experiments reveal pronounced cellular plasticity and demonstrate that the consequences of TF perturbations are highly context-dependent in tumors. Importantly, tumor cells primed for Group 3 identity, such as HDMB03 cells biased toward MYC/PR_t_ states, can be redirected toward a Group 4-like identity (Precursor/ UBC_t_) through forced expression of Group 4-specific TFs such as *EOMES*.

### PAX6 expression is associated with bi-lineage PR_t_-UBC_t_ identity in subtype VII tumors

To investigate additional molecular drivers of Group 3/4 tumor identities, we extrapolated TF-GRNs from our single-cell multi-omics atlas to a larger bulk-RNA-seq dataset of Group 3/4 medulloblastoma samples (n=703)^8,10-14^. tSNE analysis based on TF-GRN enrichment scores revealed that Group 3/4 tumors can be divided into four major axes correlating with enrichment of specific axial signatures (Fig. 5A).

**Fig. 5.**
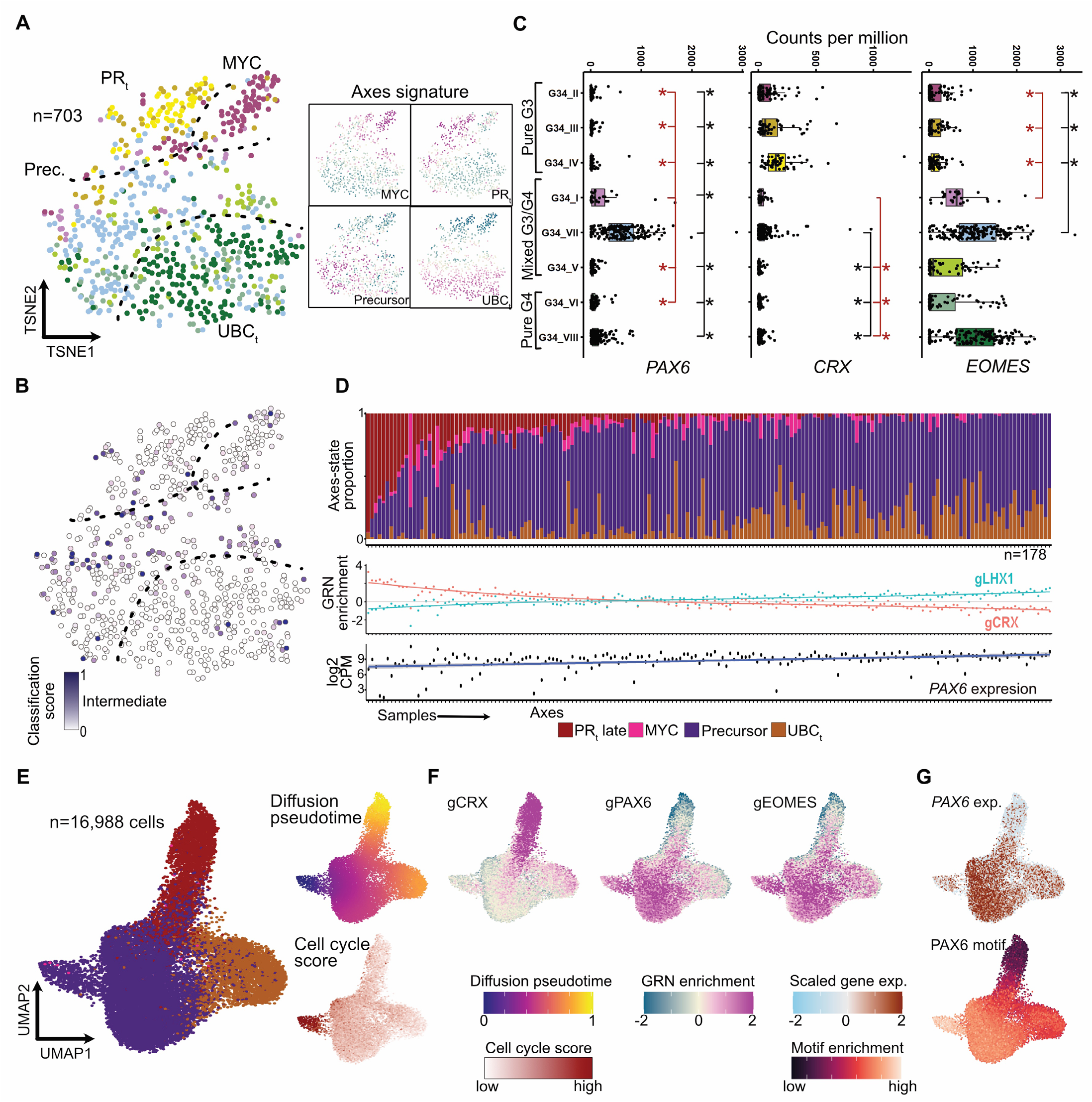
*PAX6* expression is associated with dual Group 3- and Group 4-like trajectories in Group 3/4 tumors. **A)** tSNE distribution of Group 3/4 medulloblastoma bulk RNA-seq samples (n=703) on the TF-GRN enrichment space. Relative enrichment of axial signatures (middle) and marker TF-GRNs (right) on the tSNE landscape. PR_t_, Photoreceptor (tumor)-like. Prec., Precursor. UBC_t_, UBC (tumor)-like. **B)** Intermediate methylation classification score (1-abs(G3 score-G4 score)) on the tSNE map. Dashed lines, presumptive separation among bulk axes. **C)** Boxplot distribution of *PAX6, CRX*, and *EOMES* expression in bulk RNA-seq samples (n=703) across subtypes. Expression in individual samples is shown as dots. Asterisk denotes log-fold change > 1 and adjusted p-value < 0.001 for pairwise comparisons. Black, pair-wise comparisons to subtype VII tumors, Red, pair-wise comparisons to subtype I tumors. **D)** Predicted deconvoluted axial cell-state identity in subtype VII samples arranged in order of increasing UBC_t_ identity (increasing gLHX-gCRX score). Each bar represents a sample’s proportional composition of tumor cell-states after removing predicted normal neuronal cell fraction. Middle panel, gCRX and gLHX1 AUC scores per sample. Bottom panel, *PAX6* (log2 counts per million) expression in each sample. Fitted linear model for *PAX6* expression along the sample order: R^2^=0.1598. p-value =3.241e-08. **E)** UMAP distribution of a subtype VII tumor (MB129) cells in the TF-GRN space, annotated by axial identities. Panels on right show predicted diffusion pseudotime (top) and cell-cycle score (bottom). **F)** Scaled enrichment of marker TF-GRNs in MB129 tumor cells is shown on the UMAP. **G)** UMAP distribution of scaled *PAX6* expression (top) and scaled *PAX6* motif enrichment (bottom) in MB129 tumor cells.

Overlaying common genetic driver events^8,14^ onto this landscape suggested a causal relationship between driver alterations and tumor phenotype. Predominantly *MYC*-driven subtype II tumors, defined by *MYC* amplification or *PVT1-MYC* fusion (Fig. S7A) and high *MYC* expression (Fig. S7B), were enriched for MYC and early PR_t_ axial signatures (Fig. 5A). *SNCAIP* duplication associated with *PRDM6* activation^14^ in subtypes VI, VII and VIII drove tumors towards the UBC_t_ axis (Fig. S7A).

We next focused on intermediate Group 3/4 tumors, primarily subtypes I, V and VII, which exhibited lower Group 3/Group 4 classification scores (Fig. 5B; Fig. S7C,D). In the integrated multi-omics atlas, subtype VII tumor cells were distributed along the PR_t_-to-UBC_t_ axis. Consistent with this finding, bulk RNA-seq data showed that approximately 31% (55/178) of subtype VII tumors exhibited co-enrichment of PR_t_ and UBC_t_ TF-GRN programs, and approximately 7% (12/178) showed predominance of PR_t_ TF-GRN programs (Fig. S7E). In bulk tumors, *PAX6* - a key regulator of both retinal and UBC lineage specification and differentiation^27,28^ - was significantly more highly expressed in subtype VII tumors than in other subtypes (Fig. 5C; Table S7; Table S8). Subtype VII tumors also expressed the Group 3-associated TFs *CRX* (Fig. 5C) and *NRL* (Fig. S7F) at significantly higher levels than Group 4 subtypes (VI and VIII), and the Group 4-associated TFs *EOMES* (Fig. 5C) and *LMX1A* (Fig. S7E) at significantly higher levels than pure Group 3 subtypes (II, III and IV). The intermediate identity of subtype VII therefore likely reflects co-expression of TFs from core regulatory networks governing both PR_t_ and UBC_t_ tumor cell specification. Higher *PAX6* expression also positively correlated with a greater proportion of Precursor and UBC_t_ cell states, suggesting that *PAX6* shifts tumor identity from the PR_t_ axis toward the UBC_t_ axis (Fig. 5D; Fig. S7G-K).

Genetic aberrations that could explain sustained subtype-specific *PAX6* expression, such as small variants, copy number aberrations or structural variants, have not been identified to date. Therefore, we searched for potential somatic mechanisms underlying this aberrant expression, using bulk RNA-seq data from subtype VII tumors in the ICGC cohort^11-14^. We identified a previously unreported non-coding transcript downstream of the *PAX6* locus, antisense to the *ELP4* gene (termed *ELP4-AS*; Fig. S8A; Table S9), whose expression positively correlated with *PAX6* expression (Fig. S8B; 12/21 of subtype VII samples and 1/4 of subtype I samples). We also identified samples in which *ELP4-AS* was spliced to the downstream *IMMP1L* gene (*ELP4-AS:IMMP1L*, Fig. S8C), generating a putative chimeric lncRNA in approximately 57% (8/13) of *ELP4-AS-*positive cases. Enhancer^11^ and 3D chromatin structure^29^ analyses suggest that this lncRNA:*PAX6* correlation may be linked to the gain of subtype I/VII-specific *PAX6* enhancers (Fig. S8D) that drive expression of both genes. Altogether, our findings suggest that *PAX6* is associated with dual Group 3- and Group 4-like trajectories in mixed Group 3/4 tumors.

### PAX6 drives commitment to bi-lineage PR_t_-UBC_t_ trajectory in Group 3/4 tumors

We next sought to determine whether the intermediate Group 3/4 identity arises from co-expression of dual lineage factors in the same cells, or from the presence of two distinct cell lineages in separate cells within the same tumor. We focused our analysis on three subtype VII samples (MB26, MB292 and MB129; ICGC cohort), all of which exhibited distinct bi-lineage PR_t_ (gCRX) and UBC_t_ (gEOMES) trajectories arising from a common Precursor pool (gPAX6), in clonal or sub-clonal populations (Fig. 5E-G; Fig. S9A-F; Fig S10A-C). In all three samples, *PAX6* expression and PAX6 motif enrichment were high in the Precursor cells and nearly absent in PR_t_ tumor cells (Fig. 5G; Fig. S9C,F). *PAX6* expression further correlated with Precursor/UBC_t_ markers and anti-correlated with PR_t_ markers (Fig. S9G).

To experimentally test whether *PAX6* drives this bi-lineage specification, we overexpressed *PAX6* in the *MYC*-driven cell models HDMB03 and MB3W1 (Fig. 6A). *PAX6* expression alone was sufficient to upregulate both PR_t_ and UBC_t_ signatures genes, as demonstrated by bulk-RNA-seq, in both *in vitro* cell culture (Fig. S11A,B) and *in vivo* following transplantation (Fig. 6B,C). PAX6 ChIP-seq further showed direct PAX6 binding to functional CREs at the *EOMES* and *CRX* loci (Fig. 6D), suggesting that this dual-signature upregulation is likely a direct transcriptional effect. Notably, *PAX6* overexpression drove HDMB03 tumor cells toward PR_t_ or Precursor/UBC_t_ cell states, as detected by single-cell sequencing of resulting tumors (Fig. 6E-G), with enrichment of PR_t_ and Precursor/UBC_t_ marker genes and signature gene sets in distinct tumor cell clusters (Fig. 6F,G; Fig. S11C). Mutual exclusivity of CRX and EOMES expression at the single-cell level was further confirmed in MB3W1 by immunofluorescence in resulting tumors (Fig. S11D). *PAX6*-driven cell-state changes also decreased overall *MYC* expression and promoted cell-cycle exit in cells acquiring PR_t_ identity. Importantly, commitment of *PAX6*-expressing tumor cells to bi-lineage trajectories and subsequent differentiation resulted in longer tumor latency following transplantation of HDMB03 (Fig. 6H), suggesting that the favorable clinical outcomes of *PAX6*-expressing subtype VII tumors^5^ are linked to this differentiation phenotype.

**Fig. 6.**
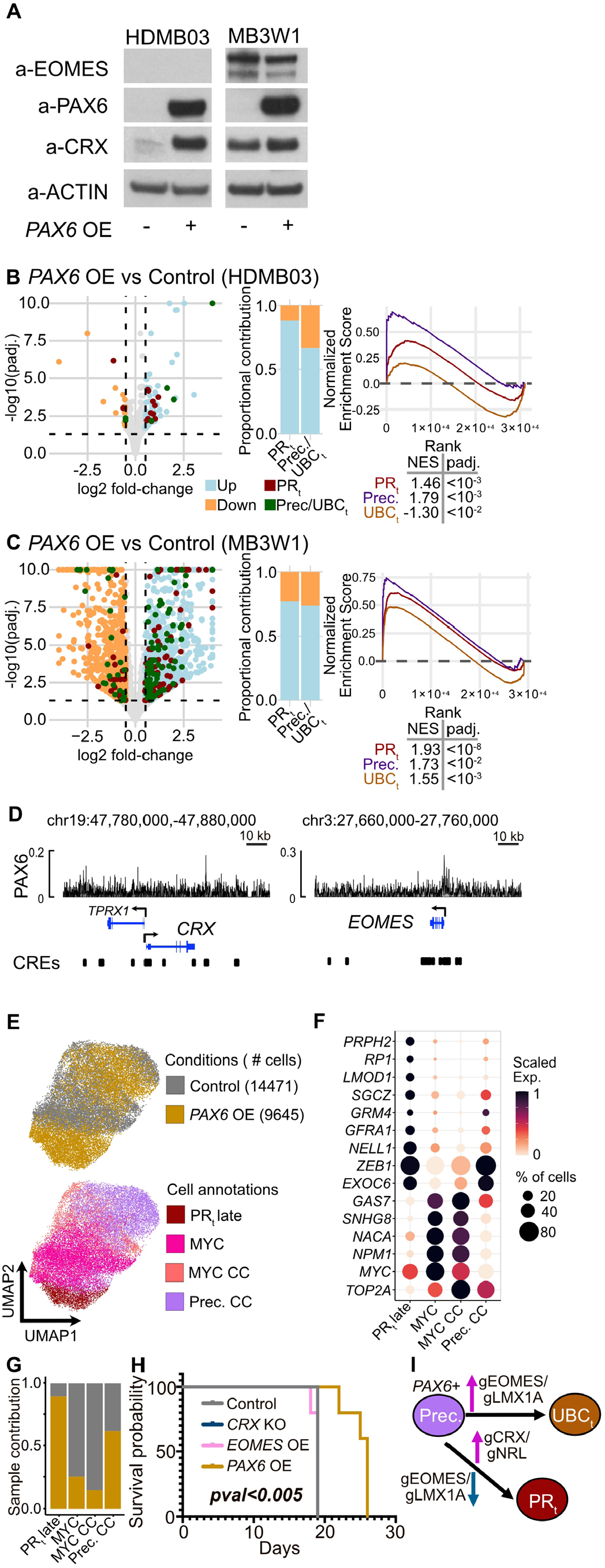
PAX6 drives commitment to bi-lineage PR_t_-UBC_t_ identity in Group 3/4 tumors. **A)** Western blot showing gain of PAX6 and subsequent CRX expression in PAX6 overexpression experiments compared to controls in HDMB03 (left) and MB3W1 (right) in vitro cell cultures. **B**,**C)** Gene expression changes in response to *PAX6* overexpression versus control in HDMB03 (B) and MB3W1 (C) *in vivo* transplanted tumors. Left: Volcano plot depicting significantly upregulated (blue dots, LFC>=0.5 and padj.<0.05) or downregulated (orange dots, LFC<=-0.5 and padj. <0.05) genes. Green dots, Group 4 gene sets. Red dots, Group 3 gene-sets. Middle: Bar plot depicting proportion of up- or down-regulated Group 3 and Group 4 signature that pass the significance thresholds (green and red dots in Volcano plot). Right: GSEA of PR_t_, Precursor and UBC_t_ gene-set. Table shows NES and padj. values for each gene-sets. **D)** ChIP-Seq signal for PAX6 in sample MB129 (this study) at *CRX* and *EOMES* loci. Bars represent CREs identified from integrated single-cell ATAC-seq. **E)** UMAP representation of scRNA-seq of HDMB03 tumor cells from control and *PAX6* overexpression. Top, cells colored by condition, number of cells per condition in parentheses. Bottom, cells colored by predicted tumor cell-states along the axial biology. **F)** Marker gene expression in tumor cell-clusters. Bubble size, proportion of cells expressing a gene per cluster. Bubble color, scaled mean value of a gene’s expression across clusters. **G)** Proportional contribution of conditions to tumor clusters. **H)** Kaplan-Meier curve of HDMB03 transplanted cells, with sacrifice occurring at first signs of tumor formation. Lines are colored by conditions. p-value, log-rank test. **I)** Axial-state transition model: Presence of *PAX6* expression leads to differentiation of tumor cells along the bifurcating PR_t_ and UBC_t_ trajectory.

Based on *PAX6* expression and motif enrichment, our experimental validation, and the established role of PAX6 in both retinal and rhombic lip development^27,28^, we propose that *PAX6* expression in the Precursor pool maintains a bi-potent state. Together with mutual repression between key PR_t_ and UBC_t_ TF-GRNs, PAX6 drives both PR_t_- and UBC_t_-identity within the same tumor in distinct tumor cell populations (Fig. 6I). The presence of divergent PR_t_/ Precursor/UBC_t_ tumor states within individual tumors thus implies a shared regulatory landscape connecting these states, and underscores that TF-GRN-driven cellular plasticity contributes to heterogeneity across Group 3/4 tumors and the intermediate identity of individual tumors.

## DISCUSSION

Despite advances in identifying a unified rhombic lip origin of Group 3/4 medulloblastoma^7-9^, the causes of heterogeneity within this group remain unknown. Our single-cell multi-omics atlas reveals the molecular underpinnings of Group 3/4 subtype-specific biology while accounting for the continuity among subtypes. Master regulators of retinal and UBC lineages, including some with key roles in both, such as OTX2, CRX, PAX6, and EOMES, are known modulators of the regulatory circuits driving Group 3/4 medulloblastoma heterogeneity^11,30^. Our analysis delineates the TF interaction network connecting these master regulators to drive divergent tumor states within a shared regulatory landscape, and proposes and experimentally validates the regulatory logic governing transitions across these states.

We show that the photoreceptor signature in Group 3/4 medulloblastoma, first reported by Kramm et al.^31^ in 1991, arises from aberrant activation of a CRX-driven photoreceptor-specification cascade, consistent with the findings of other groups^30,32^. We further show that the broad separation of Group 3 and Group 4 medulloblastoma reflects the failure of Group 3 tumors to attain *EOMES/ LMX1A-*driven UBC identity, which instead propels them toward a *CRX/NRL*-driven photoreceptor identity through remodeling of the UBC progenitor (RL_svz_) regulatory network. We propose that expression of key shared retinal factors, including OTX2 and PAX6, in UBC progenitors primes this state to acquire a divergent retinal photoreceptor identity when UBC specification stalls. Additionally, we show that beyond arising at distinct stages of UBC lineage differentiation^7-9^, the mutual repression between *CRX/NRL-* and *EOMES/LMX1A-*driven GRNs contributes to the mutual exclusion of Group 3 and Group 4 tumor identities. Finally, we provide the first single-cell resolution evidence of bi-lineage trajectories in subtype VII tumors and functionally validate the role of *PAX6* in driving this bi-lineage identity and the favorable outcome associated with this subtype.

Our results suggest that Group 3/4 tumor identity is not determined by developmental origin alone, but is instead an integrated product of cell-of-origin, oncogenic events, and regulatory mechanisms such as TF-GRNs. For example, *MYC* gain would drive tumor cells toward the undifferentiated MYC axis, while a *SNCAIP* duplication and *PRDM6* gain would drive them toward the UBC_t_ cell-state. Chromosomal copy-number variations^33^ likely similarly influence tumor cell state by modulating the activity of key TFs. We note that functional validation was performed in two MYC-driven subtype II models which, while differing in baseline expression programs, may not fully capture the plasticity of all eight subtypes. Future studies employing models representative of Group 4 subtypes will be important to extend these findings.

TF-driven lineage heterogeneity and developmental plasticity are recurrent features of pediatric brain tumors more broadly. Diffuse midline gliomas, including H3-K27M–mutant tumors, often differentiate along bi-lineage oligodendrocytic and astrocytic trajectories^34^. Diffuse hemispheric gliomas with H3G34R/V mutations, exhibit bi-lineage commitment along astrocytic and interneuron trajectories, with increased neuronal differentiation associated with favorable outcomes^35^. Other tumors display heterogeneity along more linear programs: SHH-medulloblastoma follows a granule cell lineage identity^36^; ependymomas are stalled along ependymal differentiation^37^; embryonal tumors with multilayered rosettes (ETMRs) reflect a neural lineage identity^38^; and pineoblastomas retain photoreceptor-lineage TFs, including PAX6, CRX, and OTX2, closely paralleling the photoreceptor axis described in Group 3/4 tumors^32,39^. Together, these examples illustrate that lineage-anchored transcriptional programs, whether retained, stalled, or aberrantly activated, are a common engine of intra- and inter-tumor heterogeneity across pediatric brain tumors. Beyond brain tumors, mixed-phenotype acute leukemia, including biphenotypic or bilineal subtypes, also exhibits mixed lineage trajectories^40^, suggesting a common framework across tumor types.

As with Group 3/4 medulloblastoma, such plasticity may offer a therapeutic opportunity: targeted modulation of key developmental TFs, or of signaling pathways regulated by these TFs, could force malignant cells into terminal differentiation pathways, eradicating the tumor or impeding its growth.

## Author Contributions

Conceptualization: PJ, SMP, LMK

Data acquisition: PJ, PBGdS, JN, MS, AR, CS, JS, LB, BS

Data analysis: PJ, PBGdS, TS, KO, MS, LB, MVPJ

Methodology: PJ, PBGdS, TS, KO, IS, MS, TY, KL, VH, PAN, MZ, ST

Resources: IS, MS, PS, KL, MB, JPM, KS, MBJ, PF, BJ, TM, KP, CMvT, OW, KR, AK, DTWJ

Funding acquisition: SMP, LMK

Project administration: SMP, LMK

Supervision: HK, ST, NJ, SMP, LMK

Writing – original draft: PJ, PBGdS, TS, KO, SMP, LMK

Writing – review & editing: PJ, PBGdS, TS, KO, IS, MS, TY, TM, DTWJ, VH, PAN, ST, NJ, HK, SMP, LMK.

All authors have read and approved the manuscript.

## Data and code availability

Processed snRNA-seq and snATAC-seq data generated in this study are available on GEO under the identifier GSE253557 and GSE253573, respectively. *In vitro* and *in vivo* bulk-RNA-seq and single-nuclei RNA-seq data, processed ChIP-seq summits, motif analysis are available through zenodo (17153213). Raw data will be provided after data transfer agreement. Scripts use in data processing are available on github: github.com/piyushjo15/G34MBGRN

## Acknowledgments

The study was funded in part by the European Research Council (ERC Advanced grant 819894 (Brain-Match) to S.M.P and the ERC Advanced Grant 101019268 (VerteBrain) to H.K.), the European Horizon 2020 Programme - ‘iPC - individualizedPaediatricCure’ (Grant 826121 to S.M.P). We thank the Fernando Goldsztein Family and the Cure Group 4 Consortium for their continued effort in fighting Group 4 medulloblastoma. The study was also funded in part by Fight Kids Cancer FIGHT4MB Consortium (S.M.P. and L.M.K) and the DFG Emmy Noether Program (Grant 551030459 to L.M.K.). The INFORM program is financially supported by the German Cancer Research Center (DKFZ), several German health insurance companies, the German Cancer Consortium (DKTK), the German Federal Ministry of Education and Research (BMBF), the German Federal Ministry of Health (BMG), the Ministry of Science, Research and the Arts of the State of Baden-Württemberg (MWK BW), the German Cancer Aid (DKH), the German Childhood Cancer Foundation (DKS), RTL television, the aid organization BILD hilft e.V. (Ein Herz für Kinder) and the generous private donation of the Scheu family. We also would like to express our sincere thanks to Carsten Maus, Erjia Wang (Genomics and Proteomics Core Facility, DKFZ) and Lena Weiser, Gregor Warsow (Omics IT and Data Management Core Facility, DKFZ) for their highly dedicated support in data management and processing. We also thank the flow cytometry and microscopy core facilities at DKFZ, and the DKFZ Center for Preclinical Research. We thank Kimberly Siletti for critical reading of the manuscript.

---

**Fig. S1:**
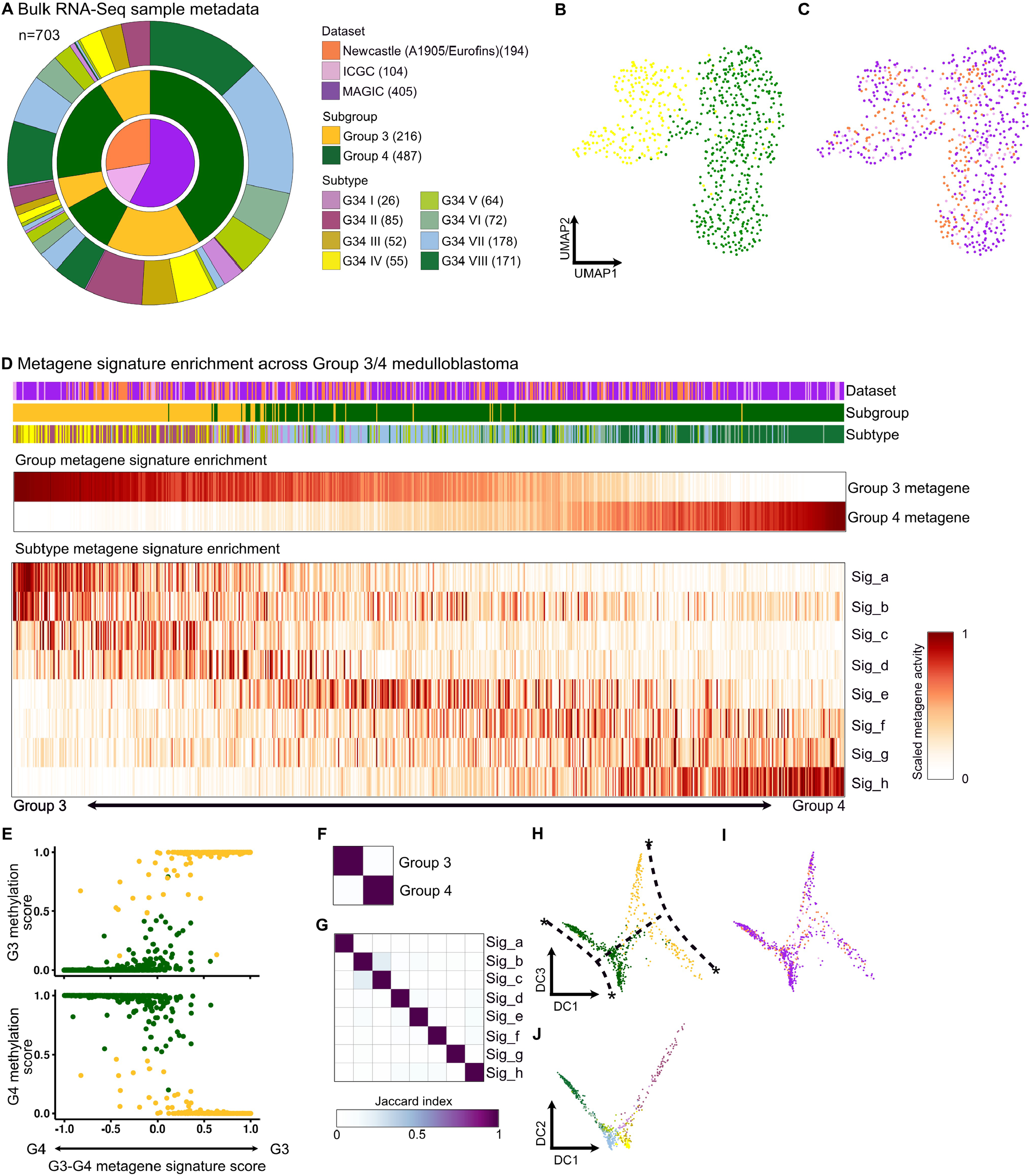
Group 3/4 medulloblastoma bulk RNA-seq data analysis and metagene signatures. **A)** Group 3/4 medulloblastoma bulk-RNA-seq metadata collated from three sources: ICGC, MAGIC and Newcastle^1-6^. Numbers of samples per category are depicted in parenthesis. **B**,**C)** UMAP distribution of tumor samples on the transcription program landscape colored by group (B) and dataset (C) identity. **D)** Scaled subgroup and subtype-specific metagene score (NMF component value) per sample. Samples are arranged on a Group 3-Group 4 metagene score scale. **E)** Per sample methylation-based Group 3 (top) and Group 4 (bottom) classification score (y axis) vs Group 3 – Group 4 metagene score (x axis). Tumor samples are colored as per subgroup identity. **F**,**G)** Jaccard similarity between subgroup-(F) and subtype- (G) specific metagene gene-sets. For each metagene gene-set, the top 100 genes ranked by contribution per metagene were used. **H**,**I)** Diffusion map of samples colored as per subgroup (H) and dataset (I) identity. **J)** Diffusion map with samples colored as per subtype identity. DC2 is shown instead of DC3.

**Fig. S2:**
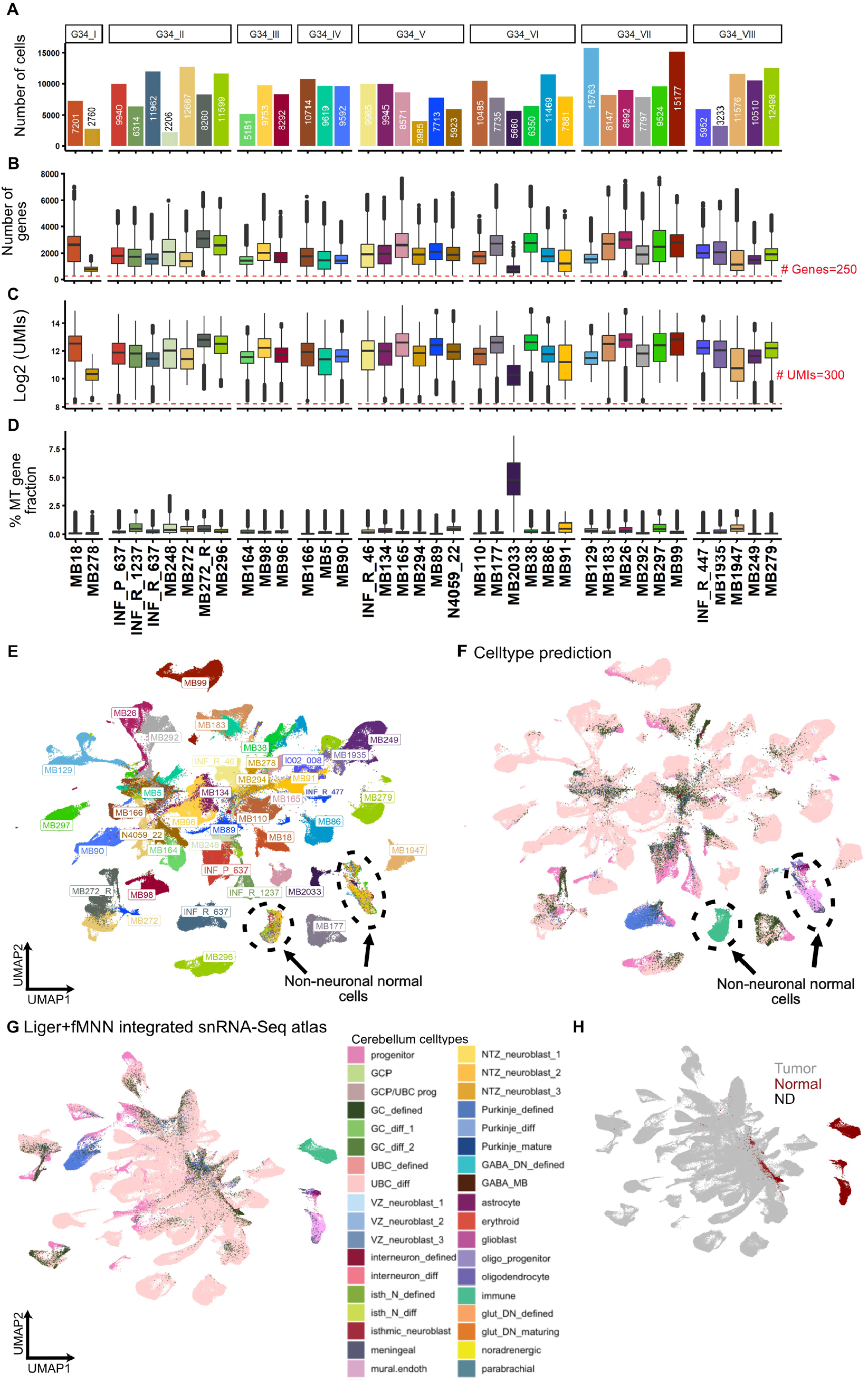
Group 3/4 tumor single-nucleus RNA-seq (snRNA-seq) data quality control (QC) metrics. **A)** Number of cells per sample in snRNA-seq data post QC filtering. **B-D**) Per sample distribution of number of genes (B), unique molecular identifiers (UMIs) (C), and fractional mitochondrial gene contribution (D). Dotted line shows cut-off at 250 Genes (B) and 300 UMIs (C). **E**,**F)** UMAP distribution of cells in the merged snRNA-seq data (without batch correction) colored by sample identity (E) and predicted cell-type labels using reference cerebellum data^7^ (F). Non-neuronal normal cells are circled. **G**,**H)** UMAP distribution of LIGER-fMNN batch-corrected snRNA-seq data colored by predicted cell-type label (G) and identified non-tumor cells (Normal and not-determined/ ND) (H).

**Fig. S3:**
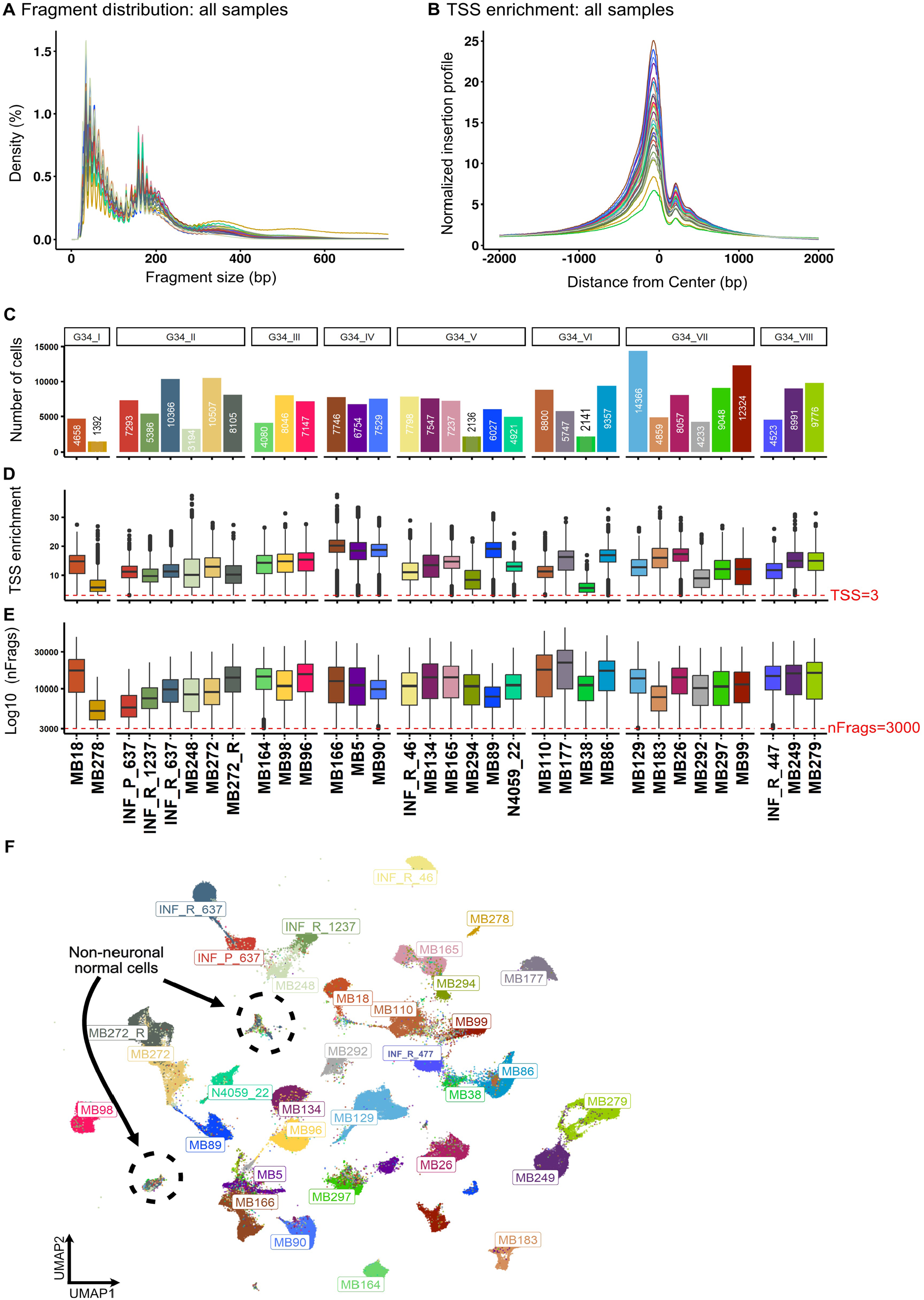
Group 3/4 tumor single-nucleus ATAC-seq (snATAC-seq) data QC metrics. **A**,**B)** Per sample Fragment size distribution (A) and Transcription Start Site (TSS) insertion profile (B). **C)** Number of cells per sample snATAC-seq data post QC filtering. **D**,**E)** Per sample distribution of TSS enrichment score (D), and number of fragments (E). Dotted line shows cut-off of a TSS enrichment of 3 (D) and 3000 fragments (E). **F)** UMAP distribution of cells in the merged snATAC-seq data (without batch correction) colored by sample identity. Non-neuronal normal cells are circled.

**Fig. S4:**
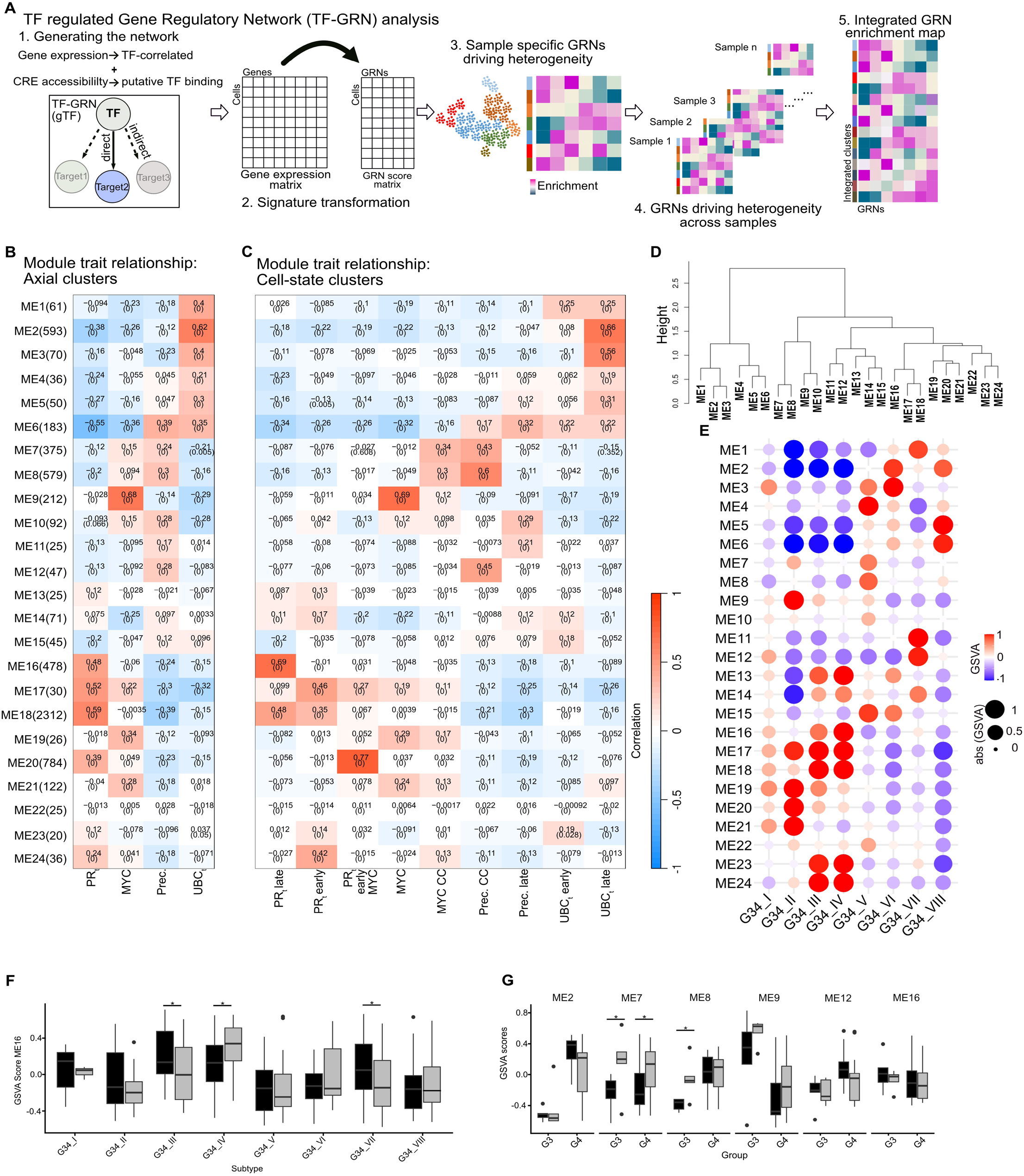
Axial gene-set signatures. **A)** Graphical representation of SCENIC+ based TF-GRN approach to integrate snRNA-seq and snATAC-seq data for the identification of the regulatory signatures driving intra-tumor heterogeneity. Conserved TF-GRNs across samples provide insights into the continuous heterogeneity observed within Group 3/4 medulloblastoma. **B**,**C)** Weighted gene co-expression network analysis (WGCNA)-based module-trait relationship correlation heatmap. Y axis are module eigengenes identified from WGCNA analysis. X axis are tumor cells clustered based on annotated axial identities (B) or cell-state identities (C). Correlation and associated p-value (in brackets) for each module-trait combination are noted in each cell. **D)** Hierarchical clustering of modules. **E)** Mean GSVA scores of module eigengenes (y axis) in tumor grouped by subtype identity (x axis). Dot color and size denotes GSVA score capped at 1 and -1. **F)** Boxplot distribution of ME16 GSVA score in primary and relapsed tumors grouped by subtype identity. Asterisk denotes p-value < 0.05 in one-sided Wilcoxon test. **G)** Boxplot distribution of selected module GSVA score in paired primary and relapsed tumors grouped by subgroup identity of primary tumor. Asterisk denotes p-value < 0.05 in one-sided paired Wilcoxon test.

**Fig. S5:**
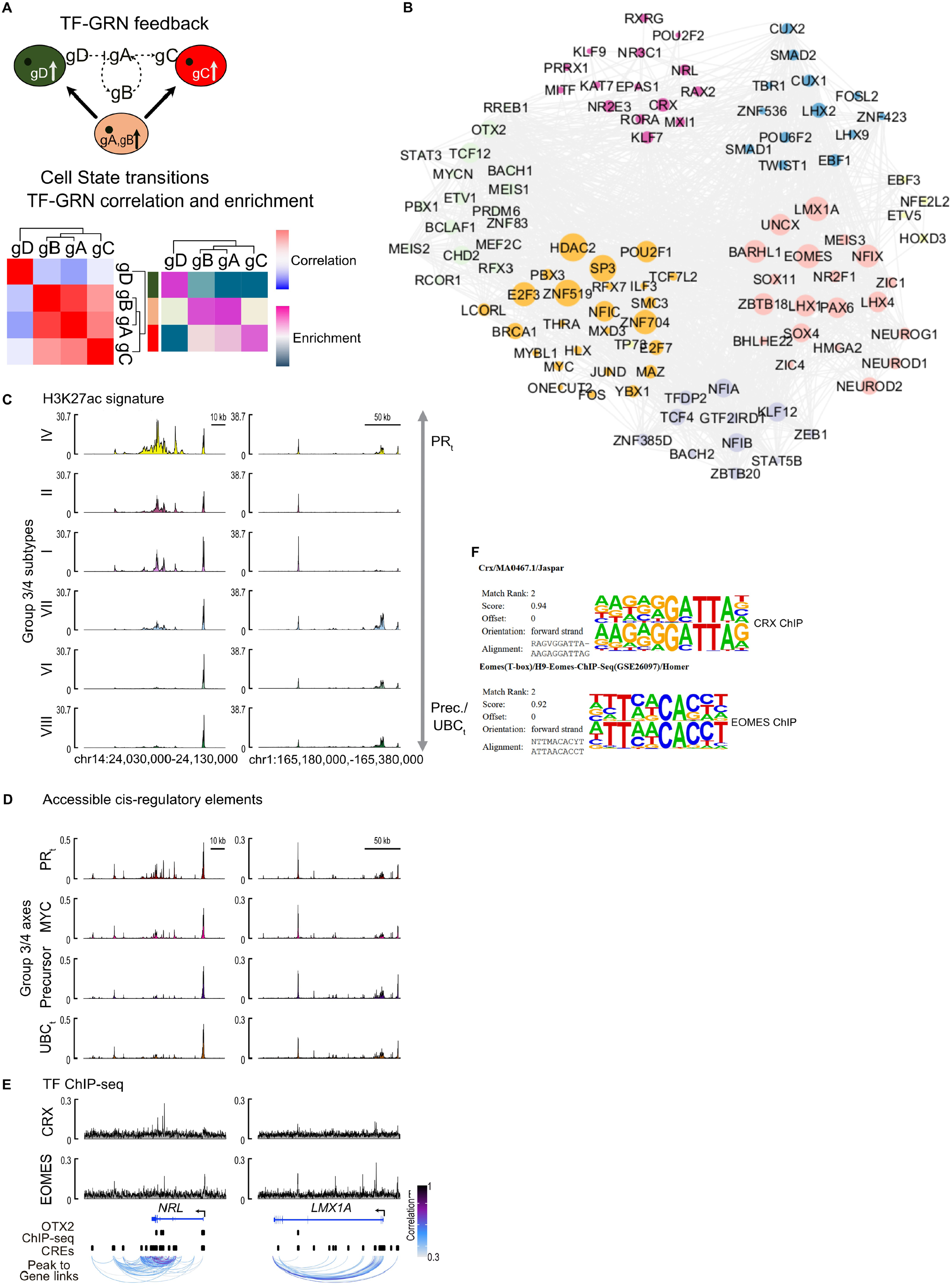
Regulatory feedback among TF-GRNs. **A)** Graphical representation of TF-GRN directed cell-state transition model. Proposed TF-GRN interactions: positive feedback loop between gA and gB. gA positively upregulates gC and gD inhibits gA. Expected correlations and enrichment of TF-GRNs per cell-state from the proposed TF-GRN network. **B)** Graph representing TF-interaction. Nodes represent individual TFs, grey lines represent interactions. Nodes are colored by TF-communities. Size of dot depicts degree of connections within a shared community. **C)** H3K27ac ChIP-seq^3^ signal profile at *NRL* (left) and *LMX1A* (right) loci in Group 3/4 medulloblastoma subtypes. Subtypes are arranged from pure high PR_t_ (top) to high Precursor (Prec.)/UBC_t_ (bottom) phenotype. **D)** Chromatin accessibility profile around *NRL* (left) and *LMX1A* (right) loci (overlapping region as selected from H3K27ac signature profile in (C)) in Group 3/4 medulloblastoma subtypes. Tumor cells were pseudobulked by axial annotation. **E)** Obtained CRX and EOMES ChIP-seq signal (this study) and OTX2 binding sites (based on published ChIP-seq data)^8^, identified CREs and representative peak-to-gene links for the selected gene (*NRL* or *LMX1A*) shown below. **F)** HOMER-based *de novo* motif identified from summit regions obtained from CRX and EOMES ChIP-seq. Identified motifs compared with known motifs.

**Fig. S6:**
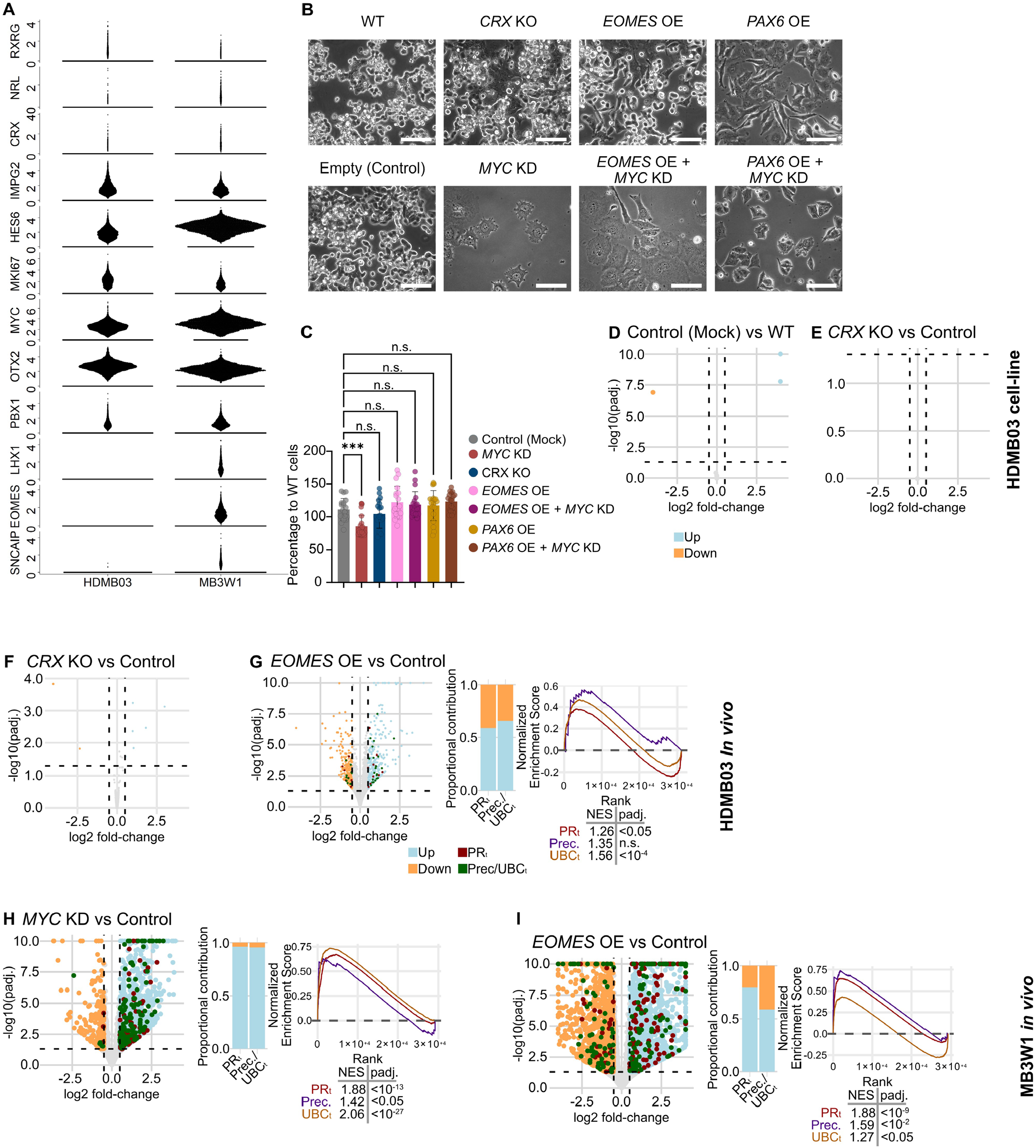
TF driven cell-state plasticity in *MYC*-driven tumor models. **A)** Beeswarm plot depicting expression of selected marker genes in HDMB03 and MB3W1 (data from GSE137758^9^). **B)** Cell morphology changes associated with altered TF expression in HDMB03 cell cultures. Scale bar, 100µm. **C)** Bar plot depicting quantification of cell proliferation changes in HDMB03 cells with control (mock) or altered TF expression, compared to WT as quantified using CellTiter-Glo. Circles, individual measurement per condition. ***, p value <0.05. n.s., not significant. **D**,**E)** Gene expression changes of HDMB03 *in vitro* experiments. Volcano plot depicting significantly expressed genes in control (mock) versus unmodified (D) and *CRX* knockout (KO) versus control (E) comparisons. **F)** Gene expression changes upon TF modulation in HDMB03 *in vivo* transplanted tumors. Volcano plot depicting significantly expressed genes in *CRX* KO versus control. **G)** Gene expression changes upon TF modulation in HDMB03 *in vivo* transplanted tumors. *EOMES* overexpression (OE) versus control. Left, Volcano plot depicting significantly upregulated (blue dots, LFC>=0.5 (right vertical dashed line) and padj.<0.05 (horizontal dashed line)) or downregulated (orange dots, left vertical dashed line, LFC<=-0.5 and padj. <0.05) genes. Green dots, Group 4 gene sets. Red dots, Group 3 gene-sets. Middle, bar plot depicting proportion of up- or down-regulated Group 3 and Group 4 signature that pass the significance thresholds (green and red dots in volcano plot). Right, GSEA of PR_t_, Precursor and UBC_t_ gene-set. Table shows NES and padj. values for each gene-sets. **H-l)** Gene expression changes upon TF modulation in MB3W1 *in vivo* transplantation experiments. Volcano plot depicting significantly expressed genes in *MYC* knockdown (KD) versus control comparisons (H) and *EOMES* OE versus control comparisons (I) *in vivo* transplantation. Left, Volcano plot depicting significantly upregulated (blue dots, LFC>=0.5 (right vertical dashed line) and padj.<0.05 (horizontal dashed line) or downregulated (orange dots, left vertical dashed line, LFC<=-0.5 and padj. <0.05) genes. Green dots, Group 4 gene sets. Red dots, Group 3 gene-sets. Middle, Bar plot depicting proportion of up- or down-regulated Group 3 and Group 4 signature that pass the significance thresholds (green and red dots in Volcano plot). Right, GSEA of PR_t_, Precursor and UBC_t_ gene-set. Table shows NES and padj. values for each gene-sets.

**Fig. S7:**
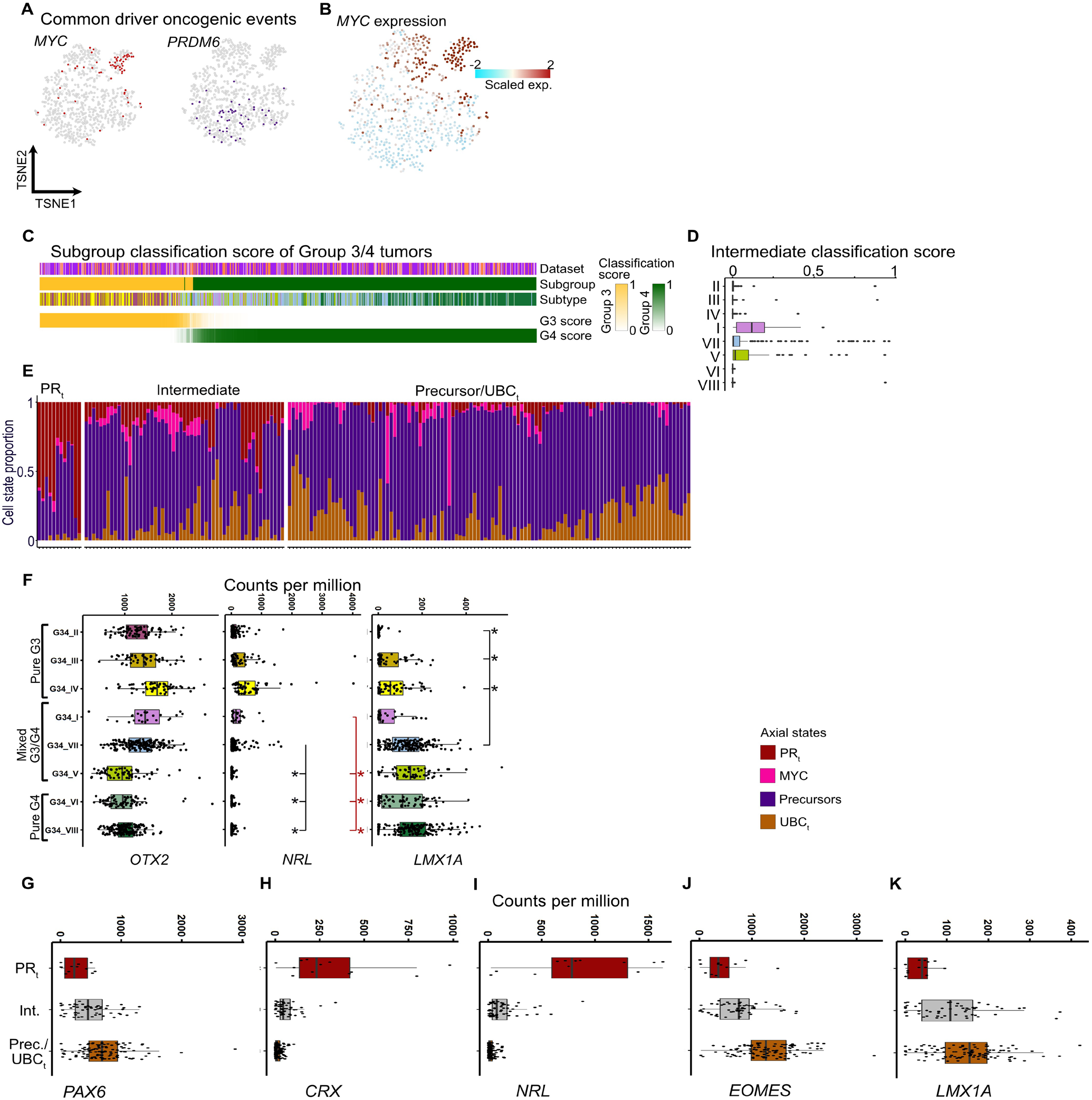
Increased *PAX6* expression is positively correlated with the UBC_t_ state in subtype VII tumors. **A)** Distribution of samples with documented genomic alterations in MYC and PRDM6 on the tSNE landscape. **B)** Scaled expression of *MYC* on the tSNE landscape. **C)** Group 3/4 tumor arranged on a Group 3 – Group 4 methylation classification score. Methylation classification score for each subgroup is on a scale of 0-1. **D)** Boxplot distribution of intermediate classification score (1-abs(G3 score – G4 score)). Samples are grouped by subtype identity. **E)** Predicted composition of axial cell-states in subtype VII samples (as shown in Fig. 5d) split by PR_t_, Intermediate or Precursor/UBC_t_ annotation. **F)**. Expression of *OTX2* (left), *NRL* (middle) and *LMX1A* (right) in bulk tumor samples across eight subtypes. Statistically significant upregulation of genes is shown by dashed lines and asterisk (log-fold change >0.5 and adjusted p-value < 0.01). Black, subtype VII tumors compared with II-IV tumors. Red, subtype I tumors compared with II-IV tumors. **G-K)** Expression distribution of *PAX6* (G), *CRX* (H), *NRL* (I), *EOMES* (J) and *LMX1A* (K) in PR_t_, Intermediate and Precursor (Prec.)/UBC_t_ annotated subtype samples, as in (E). Dots represent individual samples.

**Fig. S8:**
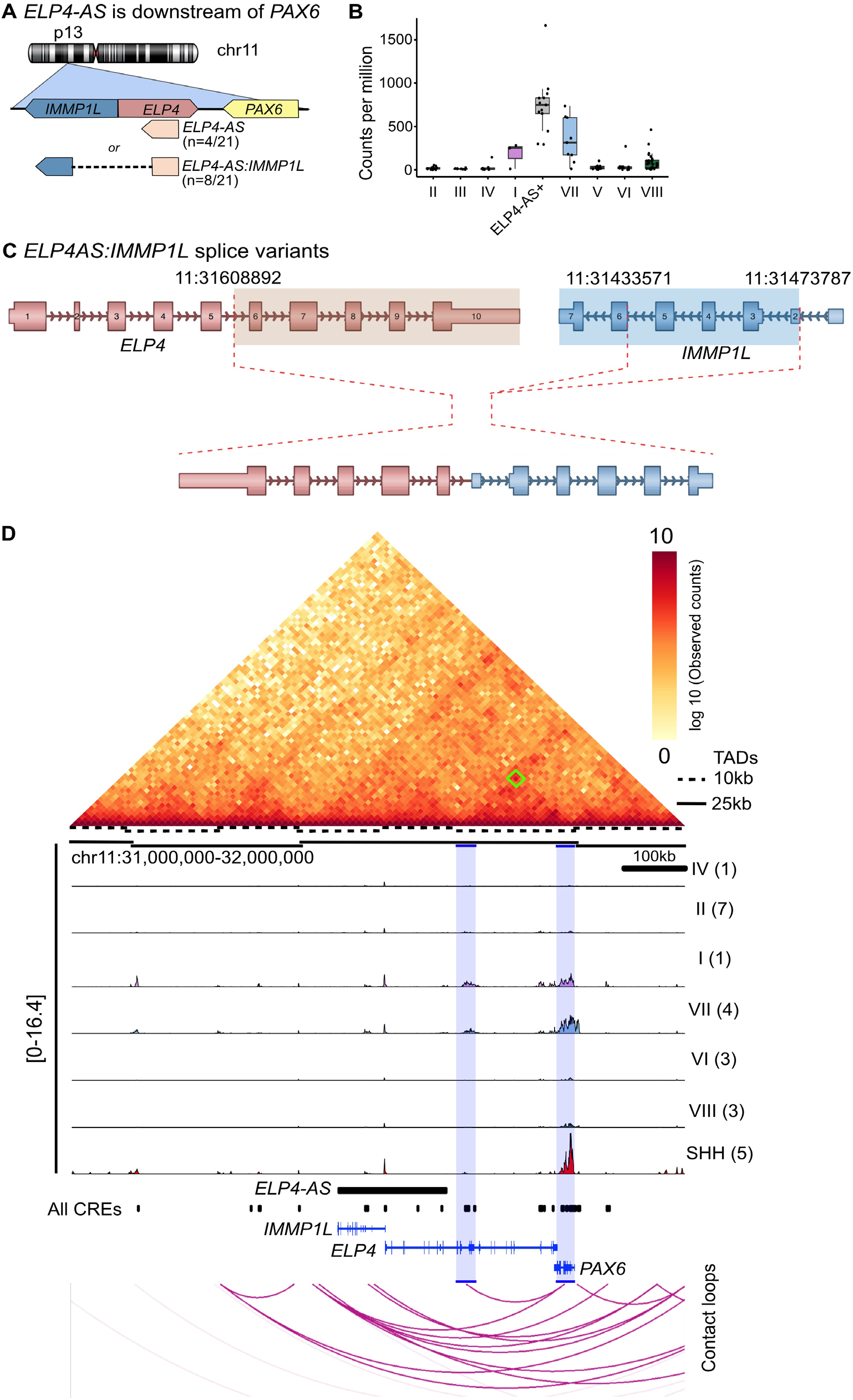
*ELP4-AS* expression is associated with *PAX6* expression in subtype VII tumors. **A)** Summary figure depicting splice events between *ELP4-AS* and *IMMP1L*. Frequency of novel lncRNA (*ELP4-AS*) and its spliced form (*ELP4-AS:IMMP1L*) in subtype VII tumors of the ICGC cohort is shown in parenthesis. **B)** Boxplot distribution of *PAX6* expression in Group 3/4 tumors (ICGC cohort). Samples with *ELP4-AS* transcripts (with or without splicing to IMMP1L, n= 13/104) (grey box) are grouped separately from the rest of the tumors, which are grouped as per subtype identity. **C)** Summary of breakpoints involved in *ELP4-AS:IMMP1L* fusion transcripts. HG38 coordinates of frequent breakpoints are shown. **D)** Top: Heatmap depicting HiC map for sample MB288^10^. Dashed and solid lines depict topologically associated domains (TADs) identified at 10kb and 25kb resolution, respectively. Middle: Trace plot shows average H3K27ac signal per Group 3/4 subtype and for SHH medulloblastoma^3^. Sample number shown in parenthesis. Bottom: Contact loops show contact points between genomic region around *PAX6* locus. Region highlighted in blue is possible chromatin interaction between subtype I/VII specific enhancer with *PAX6* loci. Green box on the heatmap denotes the contact between the blue highlighted regions.

**Fig. S9:**
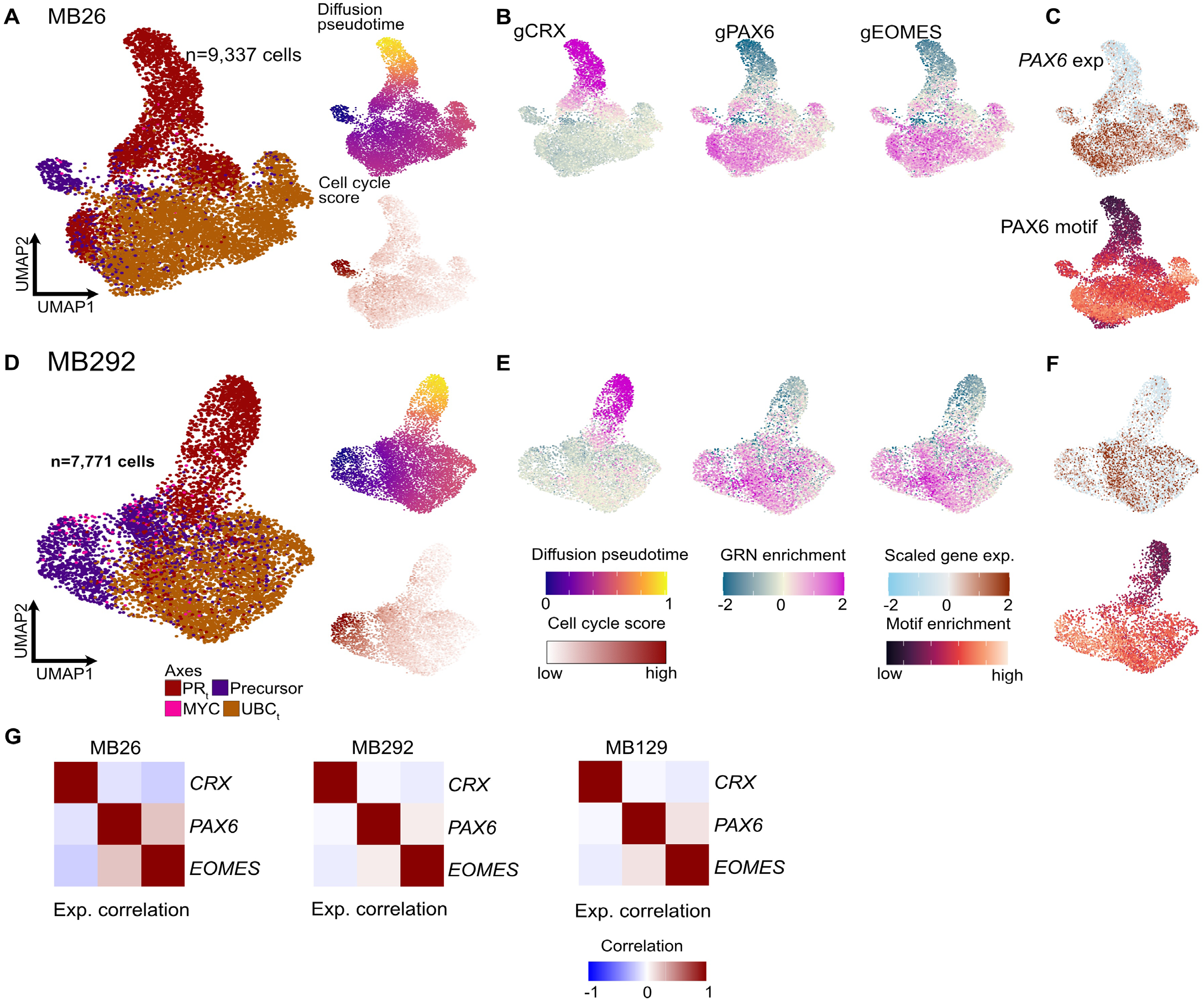
Intermediate nature of *PAX6+* subtype VII samples. **A)** UMAP distribution of tumor cells for MB26. Cells are colored as per axial identities. Panels on the left shows diffusion pseudotime (top) and cell cycle score (bottom). **B)** Enrichment of TF-GRNs gCRX, gPAX6 (Precursor) and gEOMES shown on the MB26 UMAP. **C)** Scaled PAX6 expression (top) and PAX6 motif enrichment (bottom) on the MB26 UMAP. **D)** UMAP distribution of tumor cells for MB292. Cells are colored as per axial identities. Panels show diffusion pseudotime (top) and cell cycle score (bottom). **E)** Enrichment of marker TF-GRNs shown on the MB292 UMAP. **F)** Scaled PAX6 expression (top) and PAX6 motif enrichment (bottom) on the MB292 UMAP. **G)** Scaled expression of CRX and EOMES along with their Pearson correlation with PAX6 in MB26 (left), MB292 (middle) and MB129 (right).

**Fig. S10:**
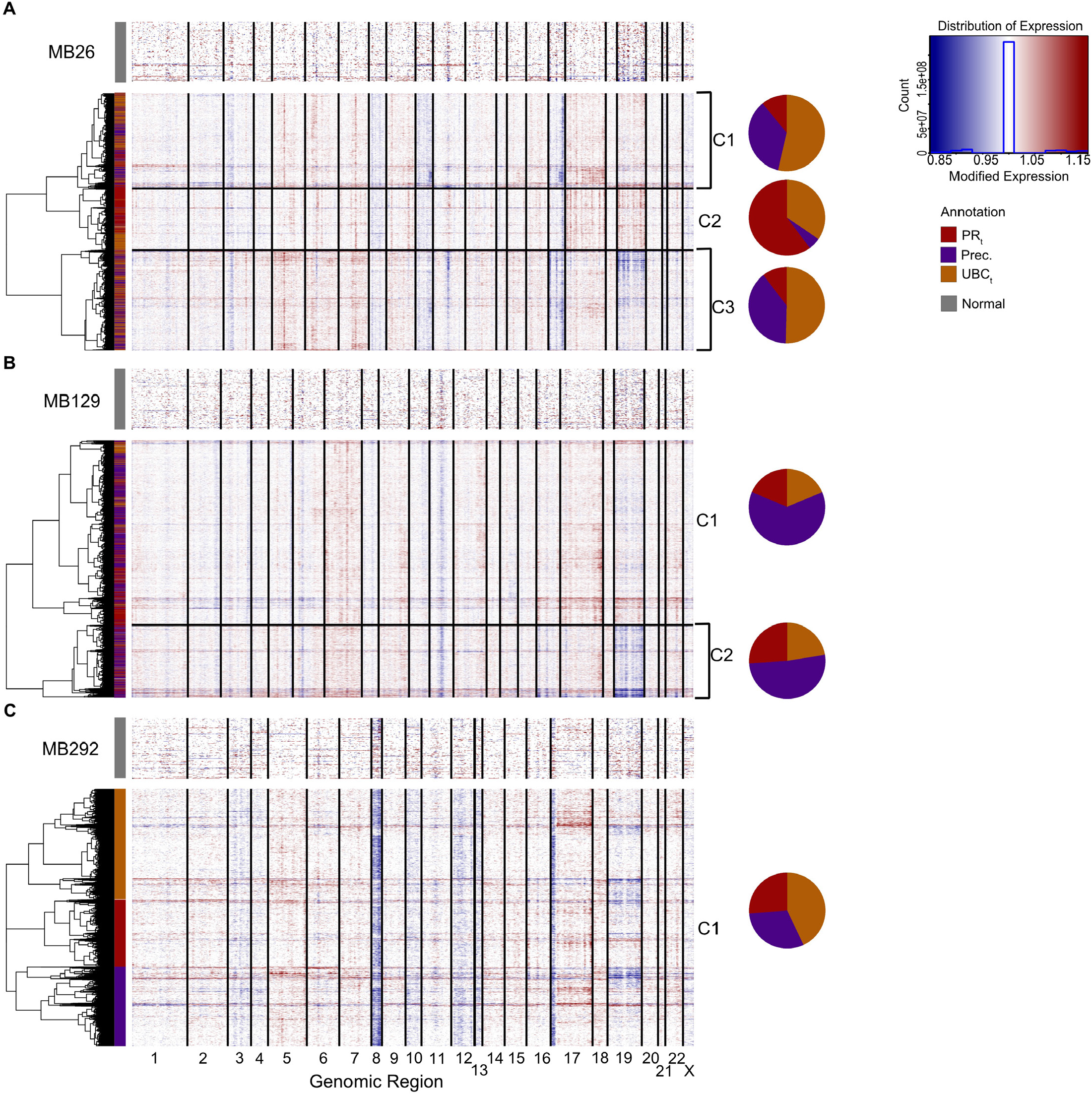
Inferred sub-clonality from snATAC-seq-derived chromosomal copy number profiles. **A-C**) Copy number variation (CNV) profiles inferred from snATAC-seq data of MB26 (A), MB129 (B), and MB292 (C). CNV analysis for MB129 and MB292 is also shown in ^11^. Three sub-clones are identified for MB26 and two for MB129. MB292 seems to be clonal. Normal reference per sample is shown on the top of each panel. Rows are colored by tumor-cell axial identities. Cells are clustered by inferred CNV signal. Left: Pie charts show axial-state composition of each subclone per sample.

**Fig. S11:**
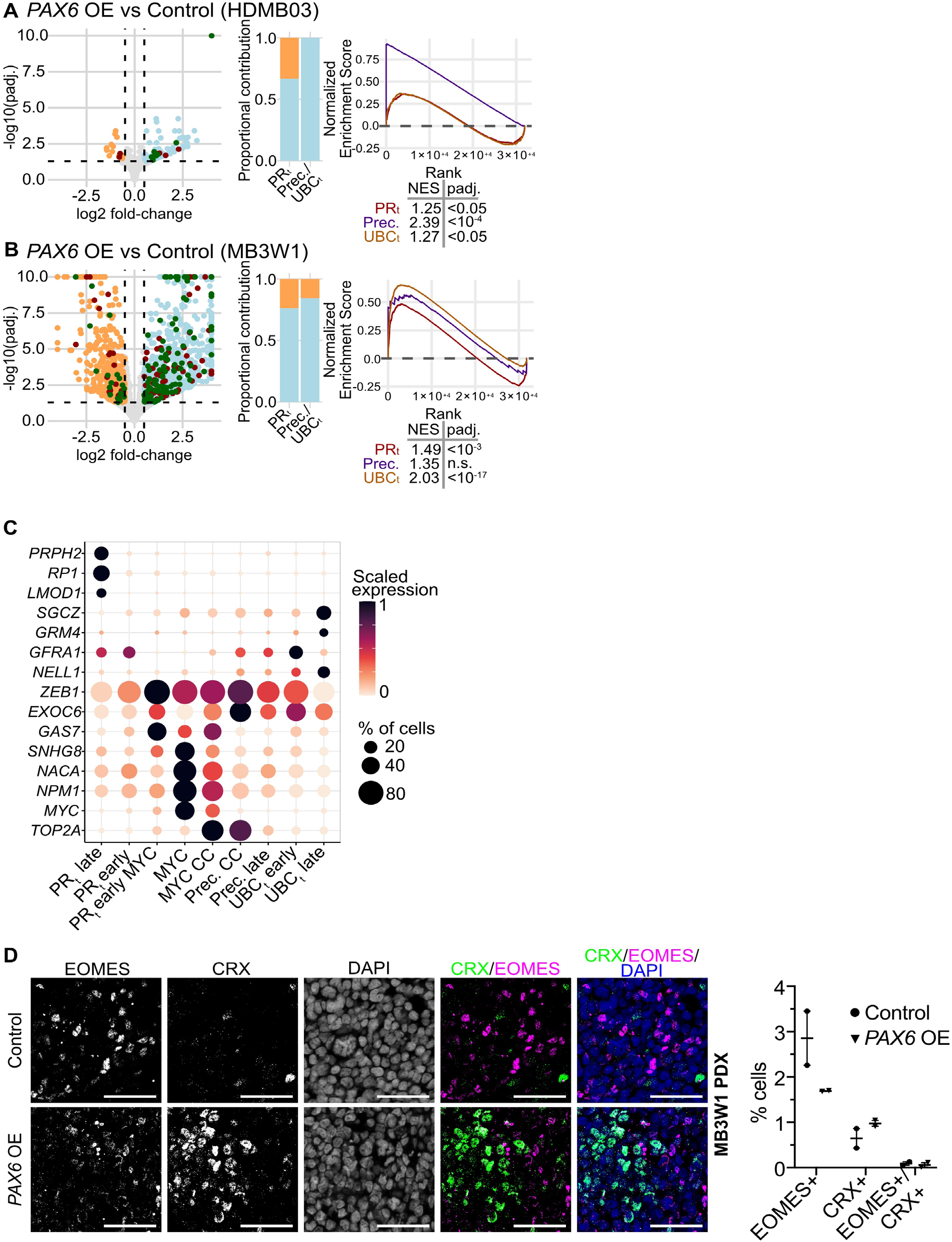
*PAX6* drives tumor differentiation toward dual Group 3- and Group 4-like cell-states. **A**,**B)** Gene expression changes in response to *PAX6* overexpression versus control in HDMB03 (a) and MB3W1 (b) *in vitro* experiments. Left: Volcano plot depicting significantly upregulated (blue dots, LFC>=0.5 (right vertical dashed line) and padj.<0.05 (horizontal dashed line)) or downregulated (orange dots, left vertical dashed line, LFC<=-0.5 and padj. <0.05) genes. Green dots: Group 4 gene sets, Red dots: Group 3 gene-sets. Middle: Bar plot depicting proportion of up- or down-regulated Group 3 and Group 4 signature that pass the significance thresholds (green and red dots in volcano plot). Right: GSEA of PR_t_, Precursor and UBC_t_ gene-set. Table shows NES and padj. values for each gene-sets. **C)** Marker gene expression in tumor cell-states in Group 3/4 tumor atlas. Bubble size depicts proportion of cells expressing a gene per cluster and bubble color depicts scaled mean value of a gene’s expression across clusters. **D)** Immunofluorescence staining marking cells expressing EOMES (purple) or CRX (green) proteins in control versus *PAX6* overexpressing transplanted MB3W1 tumors. Whisker plot (right) shows quantification of CRX+, EOMES+ and CRX+/EOMES+ nuclei (DAPI) in control and *PAX6* overexpressing tumor sections.

## Supplemental Table Legends

**Table S1**. Metadata for samples included in the bulk RNA-seq data.

**Table S2**. Metagene sets from NMF analysis of bulk RNA-seq samples. Top 100 genes ranked by contribution are shown.

**Table S3**. Metadata for samples included in the single-nucleus multi-omic atlas.

**Table S4**. 108 TF-GRN sets obtained from integrated tumor data analysis.

**Table S5**. Clustering of TF-GRNs based on co-enrichment or network graph analysis.

**Table S6**. Gene module sets obtained from the weighted gene co-expression network analysis (WGCNA).

**Table S7**. Log-fold change and adjusted p-value for selected genes in subtype I/VII pair-wise comparisons.

**Table S8**. Top 250 differentially expressed genes in subtype I and subtype VII tumors in pair-wise comparisons.

**Table S9**. ICGC samples with *ELP4-AS* or *ELP4-AS:IMMP1L* transcripts.

## Materials and Methods

### Sample selection

Target tumor tissue samples were collected from published studies (ICGC^14^ and INFORM^41^ cohorts). This study uses a portion of the cohort published in Okonechnikov et al^42^. No statistical methods were used to pre-determine the sample size. Experiments were not randomized, and investigators were not blinded to tumor sample characteristics and experiment outcome.

### Single-nucleus multi-omics sequencing

Flash frozen tumor samples were processed to extract nuclei as described^19^. Tumor samples were finely cut into pieces using a surgical blade on dry ice. Cut tissue was homogenized in the homogenization buffer (for details of reagents^19^) by trituration or douncing with a micropestle. Cellular debris was removed by centrifugation at 100g for 1 min, followed by nuclei pelleting from the supernatant at 500g for 5 min. Pelleted nuclei were washed once in the homogenization buffer before pelleting again at 500g for 5 min. Washed nuclei were re-suspended in 1x Nuclei buffer (10x Genomics) and filtered through a 40µm filter to remove the left-over debris. Nuclei concentration was estimated by counting nuclei on Countess II FL Automated Cell Counter (Thermo Fisher Scientific) with Hoechst DNA dye and propidium iodide for nuclei staining. Extracted nuclei were processed using Chromium Single Cell Multiome ATAC + Gene expression kit and Chromium Controller instrument (10x Genomics) as per manufacturer’s recommendations. One sample, MB248, was processed with Chromium Next GEM Single Cell 3’ v3.1 and ATAC v1.1 kits, as per manufacturer’s recommendation. 15,000-20,000 nuclei were loaded per channel along with the Multiome/3’/ATAC gel bead. DNA and cDNA libraries were prepared as described in respective kit protocols. Libraries were quantified using Qubit Fluorometer (Thermo Fisher Scientific) and profiled using Fragment Analyzer. GEX and ATAC libraries were sequenced using NextSeq2000 to recommended lengths and depth. If the ATAC library was not of good quality, we still used the obtained RNA-seq library if that was found to be of sufficient quality. RNA-seq and ATAC-seq datasets were further analyzed separately.

### Single-nucleus RNA sequencing (snRNA-seq) data processing

De-multiplexed reads were aligned to human genome assembly GRCh38 (v. p13, release 37, gencodegenes. org). Genome version associated comprehensive gene annotation (PRI) was customized by filtering to transcripts with the following biotypes: protein coding, lncRNA, IG and TR gene and pseudogene as recommended by *cellranger mkgtf* wrapper. Reads were aligned using STARsolo^43^ with parameters: --soloType *CB_UMI_ Simple* --soloFeatures *Gene GeneFull* --soloUMIfiltering *MultiGeneUMI* --soloCBmatchWLtype *1MM_multi_ pseudocounts* --soloCellFilter *None* --outSAMmultNmax *1* --limitSjdbInsertNsj *1500000*. For overlapping genes where intronic alignment recovered low counts, exonic alignment counts were used. Predicted cells were separated from debris using *diem* pipeline (*R*)^44^. Cells with mitochondria fraction > 1 standard deviation above the mean or above 2% (whichever is greater), and number of detected genes greater than 6600 were filtered out. We further removed cells with an intronic fraction (number of reads aligned to intron/total number of reads aligned to exon+intron) less than 25%. Filtered cells were then corrected for background signature using *SoupX* (*R*)^45^ and *celda* (*decontXcounts(), R*)^46^ pipeline. Finally, putative doublets identified by *scrublet* (*Python*)^47^ for snRNA-seq data and those identified from snATAC-seq data (see below, *Single-nucleus ATAC sequencing data processing*) were removed. Filtered gene expression matrices were normalized using the *scran* (*R*)^48^ approach. A list of 1,500 highly variable genes (HVG) per sample was also obtained after removing mitochondrial (prefix: MT-) and ribosomal genes (prefixes: RPS, RPL, MRPS, MRPL). HVG from all the samples were combined, and sex-chromosome-specific genes (chr X and Y) were further removed to obtain a set of combined sample HVG gene-sets for the single-cell cohort. Post-identification of “normal” cells (described below, *Single-cell annotation*), a list of 1,500 HVG was re-calculated from each sample and a combined tumor HVG gene-set was obtained from their union after filter sex chromosome specific genes.

### Tumor single-cell annotation

We used a published single-nucleus developing human cerebellum atlas^19^ as a reference to identify putative cell identities of each tumor cell, particularly to identify non-tumor cells, such as endothelial, immune or glial cell types. Normalized gene expression matrices from reference and target (tumor samples) were subsetted to the intersection of HVGs (5,000 genes from reference, combined sample HVG from single-cell tumor data) and cosine scaled (*cosineNorm(), batchelor, R*)^49^. A *LinearSVC* model (*sklearn. svm, Python*) was first calibrated using *CalibratedClassi fierCV(method=‘isotonic’)* (*sklearn*.*calibration, Python*) using the reference data and then the fitted model was used to assign best matching cell identities to tumor cells. Cells that were identified as immune, mural/endothelial, astrocytes or oligodendrocytes were assigned as “normal” cells. Additionally, cells identified as cerebellar granule neurons (GC-defined) but appeared as a distant cluster on UMAP, separated from the bulk of tumor cells, were also assigned as “normal”. These normal cells were removed for the integrated tumor data analysis.

### Integration of snRNA-seq data

We integrated all tumor samples together with and without batch correction (across tumor samples) using LIGER (*R*)^50^. Normalized gene-expression matrices from individual samples were subsetted to the combined sample HVG set, followed by cosine scaling. The scaled expression matrices were then used as an input for integrative NMF factorization using the function optimizeALS*(k=50, max. iters=100000)*. The obtained factors were then batch corrected using the fastMNN approach (*reducedMNN(), batchelor, R*). Corrected and uncorrected factors were used to obtain UMAP embedding of the integrated snRNA-seq data. The batch corrected factors were further used to cluster cells using KNN (*sklearn*.*neighbors, kneighbors_graph(n_neighbors=11, metric=‘cosine’, include_self=True), Python*) and leiden clustering (*leidenalg, lfind_partition*(), *Python*).

### Single-nucleus ATAC sequencing (snATAC-seq) data processing

ATAC-seq reads were aligned to GRCh38 using Cellranger’s *cellranger arc* wrapper and processed downstream using *ArchR* (*R*)^51^. Briefly, fragment files obtained post alignment were converted into arrow files (*createArrowFiles()*) using custom gene annotation (same annotation as used for snRNA-seq analysis) with a cut-off Transcription Start Site (TSS) enrichment of 3 and minimum 3000 fragments per cell. Putative doublets were identified by calculating a doublet score per cell (*addDoubletScores()*) and filterRatio of 1 (*filterDoublets())*, and were removed along with doublets identified in the snRNA-seq processing. Cells with high fragment counts, 2x standard deviation above mean, were further removed. Filtered cells were then clustered and a final QC was done by removing clusters that exhibited comparatively low TSS enrichment and number of fragments per cell, along with lack of enrichment of known marker genes, obtained from the integrated snRNA-seq data analysis. Cell clusters were also assigned putative “normal” identity if they were enriched for markers for immune, mural/ endothelial, astrocyte or oligodendrocyte lineage, based on predicted gene-scores.

### Integrating snRNA-seq and snATAC-seq data

Out of the 38 samples in the single-cell cohort, 32 were obtained using the multi-omics approach, with only a single tumor sample, MB248, using snRNA-seq and snATAC-seq data from separate cells. From here onwards, we only used tumor cell data in snATAC-seq and hence any cell identified as “normal” based on snRNA-seq or snATAC-seq processing were removed. For multi-omics data, the majority of cells had both snRNA-seq and snATAC-seq data, but as per sample, snRNA-seq and snATAC-seq data was processed separately, variable number of cells were obtained that passed QC parameters in one modality (snRNA-seq or snATAC-seq) but not in the other. To maximize data for downstream processing, we did not remove these cells from either data set, snRNA-seq or snATAC-seq, but imputed the missing RNA counts (normalized logcounts) for cells in the snATAC-seq data of the same sample. Before imputing, snATAC-seq data clusters that had RNA counts for less than 50% of cells or total number of cells with RNA counts was less than 100 were removed due to lack of a proper reference in these clusters. The imputed RNA count was then obtained from a weighted sum of normalized logcounts of 5 nearest neighbors (*sklearn*.*neighbors*.*NearestNeighbors()*). For sample MB248, snRNA-seq and snATAC-seq data were integrated using *addGeneIntegrationMatrix()* (*ArchR*).

Post integration, a joint dimensionality reduction of snRNA-seq and snATAC-seq data was obtained per sample. Using *addCombinedDims()* (*ArchR*), we combined Latent Semantic Indexing (LSI) based factorization of snATAC-seq data to singular value decomposition (SVD) based factorization of snRNA-seq data, excluding dimensions that had a correlation of greater than 0.75 to sequencing depth. The joint dimensional reduction was used to identify clustering (referred to as *combined_cluster*) and UMAP representation of the combined ATAC-RNA data.

### Per sample peak calling in the snATAC-seq data

Peaks were called per sample on the tumor cells grouped by *combined_cluster* annotation. First a minimum of 40 cells and a maximum of 500 cells per group, with a sampling ratio of 0.8, were used to generate pseudobulk replicates via *addGroupCoverages()*. Then peaks were identified using MACS2 caller with a reproducibility of 2 via *addReproduciblePeakSet()*. The rest of the parameters used were defaults as defined in the function definition.

### Creating a cisTopic object per sample

To prepare data for *SCENIC+* pipeline (*Python*)^15^, the “peaks by cells” matrix (referred to as peak matrix here onwards) obtained from the *ArchR* analysis was converted to cisTopic object (*pycisTopic, Python*) to obtain topics and differentially accessible regions (DARs), which represent candidate enhancers for *SCENIC+* analysis. Peak matrix was reduced to 50 topics (*run_cgs_models(), pycisTopic*), obtained topics were binarized into region sets by ‘*otsu’* method and selection of top 3,000 regions per topic. DARs were identified by first identifying highly variable features (HVF), based on the log-normalized peak matrix, and then identifying marker regions using a cut-off adjusted p-value less than 0.05 and Log2FC greater than 0.5. If no marker regions were identified, then lower thresholds (Log2FC <0.1 and adjusted p-value <0.5) were used.

### Creating motif-enrichment dictionary

Candidate enhancer regions identified from topic analysis and DARs were then assessed for motif-enrichment leading to creation of cistromes, an object associating transcription factors (TFs) to potential target regions. We used *run_ pycisTarget()* wrapper from *SCENIC+*, along with motif-ranking, motif-score and motif-annotation provided by the Aertslab for GRCh38^15^ to obtain the TF-region cistromes per sample. Default settings were used for the function with the exception of *run_without_promoters = True*. Further, only TFs that were present in the combined HVG set were selected for further processing.

### Gene regulatory network identification

We used the *SCENIC+* approach for the multi-omics data to identify TF-associated gene regulatory networks (TF-GRNs) per sample. To identify tumor TF-GRNs, we first removed cells that were assigned as “normal” identity in snRNA-seq or snATAC-seq data processing. For each sample, we used snRNA-seq data (after converting it into anData object), snATAC-seq data (as cisTopic object) and motif-enrichment dictionary (obtained from pycisTarget) to create a *SCENIC+* object. Additionally, we provide a TF adjacency matrix with correlation values from a separate run of *pyscenic* (*Python*)^52^ using *‘genie3’* method (-m flag). *SCENIC+* first identified region-to-gene linkage for identified enhancers and their target genes and then assigned TF-to-gene links by associating TF that are enriched in the enhancers found linked to target genes. In the final step, *SCENIC+* uses region-to-gene and TF-to-gene links to identify regulons (TF-to-region-to-gene links) that are among the top ranked based on importance scores and assigns positive or negative regulatory relationships based on the correlation between the TF and assigned target gene. *SCENIC+* outputs a list of possible regulons with putative activation or repression relationships. For our analysis, we focused on positive TF-target interactions, represented as ‘+_+’ in *SCENIC+*.

### TF-GRNs selection and compilation

For each sample, a set of active TF-GRNs was identified using *SCENIC+* approach as described above. For each of the TF-GRNs, an “Area Under the Curve” (AUC)-based enrichment score (*AUCell_run(), AUCell, R*)^52^ was calculated for all the tumor cells using log normalized RNA counts (including the imputed counts). From the identified TF-GRNs, GRNs associated with heterogeneity were identified based on the differential enrichment of TF-GRN AUC scores across combined_cluster annotation using Wilcox-rank test (*findmarkers(), scran, R*). The top three marker TF-GRNs per cluster per sample were used as representative of differentially active GRNs for that sample. After identifying such sets of TF-GRNs for each sample, we combined the obtained gene-sets as follows: 1) we selected TFs that were found to be associated with differentially active TF-GRNs in at least two samples, and then 2) for each of these selected TFs, we filtered target genes that were identified as linked to the TF in more than 25% of the samples where the TF was found to be active, with the association being present in at-least two samples. TF-GRN sets with sizes of less than 15 genes (including the TF) were also removed. In this way, we identified a conserved set of TF-gene links that were biologically replicated while reducing the number of associated genes by increasing the number of replicates required for the TFs that were widely used. This resulted in 108 TF-GRNs (Supplementary Table S4).

### Integrating tumor RNA data across samples using TF-GRN enrichment scores

We obtained the AUC score for each of the TF-GRN gene-sets (n=108) for all of the tumor cells using *AUCell_ run(aucMaxRank=0*.*1*nGenes, normAUC=TRUE) (AUCell)*. The resulting score matrix was factorized using NMF (rank=25) and the obtained NMF factors were used for clustering (KNN-leiden) the integrated tumor data (resulting in 96 clusters), and obtained UMAP embedding and diffusion plots (*destiny, R*)^53^. We used *addmodulescore()* (*Seurat, R*)^54^ to calculate gene-set activity scores for each of the 108 TF-GRN sets in the combined tumor gene expression data. The TF-GRN activity score matrix was scaled across cells and averaged per cluster to obtain the TF-GRN enrichment heatmap. The scaled TF-GRN matrix (clusters x TF-GRNs) was hierarchically clustered to obtain groups of co-enriched TF-GRNs (annotated as TF-GRN programs) and groups of tumor cluster exhibiting similar TF-GRN activity (annotated as tumor axes and cell-states). The TF-GRN activity score was also used to obtain Pearson correlation between TF-GRNs.

### Integrating snATAC-seq data across samples

*ArchR* generated arrows files across tumors were merged to obtain a combined *ArchR* object. The merged *ArchR* object was factored using *addIterativeLSI(iterations=5, clusterParams = list(resolution = c(0*.*1, 0*.*2, 0*.*4, 0*.*8), sampleCells = 20000, n*.*start = 10), varFeatures = 100000, dimsToUse = 1:100, totalFeatures = 500000)* and obtained factors were used to calculate joint UMAP representation of the snATAC-seq data. The merged ArchR object was then subsetted to tumor cells to identify peaks in the integrated data. Similar to peak identification in individual samples, first the integrated data was pseudobulked by tumor cell clusters (as identified in *Integrating tumor data using TF-GRN enrichment scores*) using *addGroupCoverages(maxCells = 1000, minReplicates = 5, maxReplicates = 15, maxFragments = 50 * 10^6)*. Peaks were called using *addReproduciblePeakSet(reproducibility = “2”)*. The frequency of the identified peak’s activity per tumor cluster was calculated by dividing the number of cells in a cluster in which the peak was detected by the total cluster population. Peaks that showed less than 3% frequency in all the tumor clusters were filtered out to obtain a robust peak set.

### TF-GRN cis-regulatory elements (CREs) activity in tumor cells

Cis-regulatory elements (CREs) associated with a candidate TF and its identified target genes were combined to obtain a non-overlapping region set that defined the putative functional binding regions of that TF. For each TF-GRN, the obtained CREs were filtered to those CREs that overlapped with the above identified robust peak set (see *Integrating snATAC-seq data across samples*), which together represented a pseudo-peak for that TF-GRN. A TF-GRN x tumor cluster pseudo-peak counts matrix was obtained by summing the peak counts of the associated CREs per tumor cluster. This matrix was divided by sum of column values, scaled to 10,000, and finally log_2_ transformed to obtain a normalized CRE activity matrix. The normalized CRE activity matrix was scaled across rows to obtain the CRE enrichment heatmap.

### Weighted gene co-expression network analysis (WGCNA) analysis

We used the combined log-counts and final annotation for the tumor data to identify a set of genes that showed axes or cell-state correlated activity using WGCNA (*R*)^55^. A normalized gene expression matrix was subsetted to a combined tumor HVG set. For the WGCNA run, softPower was set to 9 and minimum module size was set to 20. A total of 24 modules were identified. To analyze enrichment of WGCNA module gene-sets per group 3/4 tumor subtypes, GSVA scores were obtained for gene-sets per sample cohort (ICGC, MDT, and Newcastle) using *gsvaParam()* and *gsva()* (GSVA,R)^56^. GSVA scores were scaled per cohort and then averaged per subtype. GSVA scores per gene-set were also obtained per sample for primary-relapse cohort Medulloblastoma, Korshunov, n=643, group 3/4 n=435), and paired primary-relapse cohort (Medulloblastoma, Korshunov, n=86, group 3/4 n=38) obtain from R2: Genomics Analysis and Visualization Platform (https://r2.amc.nl).

### Bulk RNA-seq data processing

The patient bulk-RNA-seq data was collated from published studies for three cohorts: ICGC^11-14^, MAGIC ^8^ and Newcastle^10^. Except for the ICGC cohort, processed read count matrices were used for MAGIC and Newcastle samples. For samples belonging to the ICGC cohort, raw reads were aligned to human genome assembly GRCh38 (v. p13, release 37, gencodegenes.org), using STAR aligner^57^. RNA-seq samples belonging to individual cohorts were normalized separately using *DESeq2* (*R*)^58^. Intersection of genes among the top 5,000 HVGs per cohort were used for subsetting data for NMF factorization. NMF factorization was performed using *sklearn*.*composition*.*NMF (init=“nndsvd”, max_iter=100000)*. NMF rank 2 and 8 were used to obtain subgroup- and subtype-associated latent factors or metagene signatures. To obtain the gene-set associated with each latent factor/metagene signature, the top 100 genes ranked by contribution to that factor were used. The obtained NMF latent factors (rank=8) were used for UMAP, tSNE, and Diffusion map projection of the bulk data. Differentially active genes in subtype I or VII tumors were obtained from pairwise comparison using *lfcShrink(type=“ashr”)* (*DeSeq2*).

Bulk RNA-seq data generated from HDMB03 and MB3W1 cell cultures or transplanted tumors were processed using the same pipeline as described for the ICGC cohort. Differential gene expression analysis was performed within each tumor cohort (HDMB03 or MB3W1; cell line or frozen tumor samples) using *lfcShrink(type=“ashr”)*. Volcano plots were created using *ggplot()*.

### Deconvolution of bulk RNA-seq tumor data

Bulk RNA-seq data was deconvoluted using *BayesPrism* (*R*)^59^ separately for each cohort (ICGC, MDT, Newcastle). Tumor data with cell-state annotation was combined with non-neuronal cells from single-nucleus human cerebellum data^19^ to create the reference for deconvolution. An intersection of combined single-cell multi-omics atlas derived tumor HVG set and the top 5000 HVGs from the bulk tumor cohort was used to subset the gene expression matrices of the reference and target data. Estimated proportion for each of the reference cell-state were obtained for each of the tumor sample, and combined estimate of the non-neuronal cells were removed to obtain the proportional composition of tumor cells in terms of the reference cell-states as annotated in the integrated Group 3/4 medulloblastoma atlas.

### Gene-set activity scores for bulk RNA-seq data

TF-GRN activity scores were calculated for each of the bulk tumor samples using*addmodulescore()*. Gene-set activity scores for tumor samples were scaled for each cohort (ICGC, MDT, and Newcastle) separately and then merged, and scaled scores were used to obtain tSNE representation of the bulk-RNA-seq tumor data on the TF-GRN enrichment space.

### Gene-set enrichment analysis (GSEA) for axial-state signatures in bulk RNA-seq data

Differentially expressed genes statistics were obtained per comparison using *lfcShrink(type=“ashr”)* and genes were ranked as per *foldchange sign x -log*_*10*_ *(pvalue)* in decreasing order. Representative gene-sets obtained from WGCNA analysis for PR_t_ (ME16), Precursor (ME12), and UBC_t_ (ME9) were used for gene-set enrichment analysis using *fgseaMultilevel*(*scoreType=‘std’*) (*fgsea, R*, https://github.com/alserglab/fgsea).

### TF Chromatin Immunoprecipitation sequencing (ChIP-seq)

CRX, EOMES and PAX6 ChIP library preparation was performed at Active Motif Services (Carlsbad, CA) using antibodies against CRX (#AF7085, R&D Systems), EOMES (#66325, CST) and PAX6 (#PRB-278P, Biolegend). Frozen patient tumor tissues were used to prepare chromatin: EOMES (MB165, MB129, MB297), CRX (MB96, MB129, MB297), PAX6 (MB129). Briefly, tissue was pulverized with mortar and pestle in liquid nitrogen. Pulverized tissue was then fixed in PBS + 1% formaldehyde at room temperature for 15 minutes and fixation was stopped by the addition of 0.125 M glycine to final concentration. Chromatin was isolated by adding lysis buffer and sonicating using the PIXUL® Multi-Sample Sonicator (Active Motif, Catalog #53130) to shear DNA to an average fragment size of 200–1000 bp. To determine chromatin yield, an aliquot of the sheared chromatin was reverse crosslinked at 65°C, treated with RNase and proteinase K, and subjected to DNA purification using SPRI beads (Beckman Coulter). DNA concentrations were measured using a Qubit Fluorometer (Thermo Fisher), and total chromatin yield was extrapolated based on the original chromatin volume. For ChIP reactions, aliquots of chromatin were precleared with protein G agarose beads (Invitrogen). Immunoprecipitations were performed using antibodies specific to the target proteins of interest. After washing, immune complexes were eluted from the beads using SDS buffer, treated with RNase and proteinase K, and de-crosslinked by overnight incubation at 65°C. ChIP DNA was then purified using phenol-chloroform extraction and ethanol precipitation.

ChIP DNA libraries were prepared using either the PrepX DNA Library Kit (Takara Bio) on the Apollo automation platform or the NEB DNA Library Prep Kit, following the manufacturers’ protocols. ChIP-seq libraries were sequenced with 2 × 50 bp paired-end reads on the Illumina NextSeq 2000, resulting in on an average 20 million read pairs per sample.

### ChIP-seq data analysis

Published H3K27Ac ChIP-seq data^11^ was aligned to GRCh38 using *bowtie2*^*60*^. Duplicated, unmapped and multi-mapped reads were marked and removed using *sambamba*. Deduplicated alignment bam files were sorted using *sambamba* and indexed using *samtools*. Obtained alignment was normalized using *bamCoverage –normalizeUsing CPM–binSize 20 smoothLength 60 –extendedReads 150 –bl hg38. blacklist*.*v2*.*bed* (*deepTools, Python*) and converted into bigwig format. Enhancer signal for a subtype was obtained from averaged normalized signal of the constituting samples using *wiggletools. wigToBigWig* was used to convert obtained *Wig* files to bigwig and followed by conversion to *BedGraph* format using *bigWigtoBedGraph* tool. Bed files for human OTX2 (GSE137311), binding regions were obtained from Remap (https://remap2022.univ-amu.fr/). Track plots were prepared by *SparK* (https://github.com/harbourlab/SparK). TF-ChIP-seq data was aligned to GRCh38 using *bwa*^61^. Aligned reads were processed as described above. ChIP-seq signal were visualized as described above for enhancer data. Peaks were called using *macs3 (-f BAMPE -p 0*.*001)* (https://macs3-project.github.io/MACS/). Summits obtained for a TF from all the samples were merged using *bedtools merge* to obtain representative binding regions. Motif analysis was performed using HOMER^62^ to identify *de novo* motifs.

### Identification of *ELP4-AS* and *ELP4-AS:IMMP1L*

Novel long non-coding RNA transcript, *ELP4-AS*, was identified using *StringTie* based *de novo* transcriptome inoculated mouse tissue were mechanically dissociated, passed through a 40 µm cell strainer, and seeded as single-cell suspensions in a defined medium consisting of DMEM/F-12 (Thermo Fisher), 2% B27 supplement (Thermo Fisher), 1% MEM vitamin solution (Thermo Fisher), 20 ng/mL basic fibroblast growth factor (bFGF; Preprotech), 20 ng/mL epidermal growth factor (EGF; Preprotech), and antibiotics (40 U/mL penicillin, 40 μg/mL streptomycin). Cells were sub-cultured at split ratios of 1:3 to 1:5 by mechanical dissociation.

HEK293T cells were grown as adherent monolayers in high-glucose DMEM (Thermo Fisher) supplemented with 10% FBS (Sigma-Aldrich) and antibiotics (40 U/ mL penicillin, 40 μg/mL streptomycin; Sigma-Aldrich). Cultures were passaged at 80% confluence using trypsin-EDTA (Sigma-Aldrich) at split ratios of 1:5 to 1:10.

All cultures were incubated at 37°C in a humidified assembly using the ICGC cohort RNA-seq data. The atmosphere containing 5% CO2 and the medium was spliced variant of *ELP4-AS* with downstream *IMMP1L* was identified using *Arriba* toolkit based on the RNA-seq data^12^. Presence of *ELP4-AS* and novel splicing transcript was confirmed by RT-qPCR in individual samples. Presence of fusions at genome level was also investigated using WGS data and SOPHIA algorithm^13^.

### HiC data analysis

Processed HiC data (.hic) was downloaded for the sample MB288 from GSE240985^10^. HiC heatmap was plotted using *plotHicTriangle()* (*plotgardener, R*)^14^. Contact loops were obtained using *peakachu*^*15*^ at 10kb resolution. Topologically associated domains (TADs) were obtained using *SpectralTAD* (*R*)^16^ at 10kb and 25kb resolution.

### Copy number variation (CNV) analysis in single-cell data

snATAC-seq data was used as an input to infer CNV profile per sample using *atacInferCNV*() *(https://github.com/kokonech/atacInferCNV, R)*, similar to published work^33^.

### Cell culture

HDMB03 cells were maintained as semi-adherent cultures in RPMI-1640 medium (Thermo Fisher) supplemented with 10% fetal bovine serum (FBS; Sigma-Aldrich) and antibiotics (40 U/mL penicillin, 40 μg/mL streptomycin; Sigma-Aldrich). Cells were passaged at approximately 80% confluence using trypsin-EDTA (Sigma-Aldrich) at split ratios ranging from 1:2 to 1:10.MB3W1 cells were cultured under serum-free, neurosphere-promoting conditions. Cells isolated from refreshed every 2–3 days. Cells were verified regularly as mycoplasma free.

### Molecular cloning and CRISPR–Cas9 sgRNA design

Cloning was performed using the In-Fusion® Snap Assembly Master Mix (Takara Bio) according to the manufacturer’s instructions. Coding sequences of *EOMES* (RefSeq ID: NM_005442.4) and *PAX6* (RefSeq ID: NM_000280.6) were amplified using primers designed with the Takara In-Fusion primer design tool for insertion into the pCDH-mCherry backbone vector. Each assembly reaction was performed at a 1:5 molar ratio of vector to insert and incubated overnight at 50°C. One Shot™ Stbl3™ chemically competent E. coli (Thermo Fisher) were transformed with the reaction products, and positive clones were expanded for plasmid preparation. All constructs were verified by Sanger sequencing to confirm full sequence integrity prior to use.

For CRISPR-Cas9 knockout experiments, single guide RNAs (sgRNAs) were designed using the ChopChop online platform. Four sgRNA candidates against *CRX* were selected, cloned in px330 plasmid (Addgene #42230) and functionally tested using the pCAG-EGxxFP reporter assay (500 bp targeting sequence was cloned using In-Fusion as previously described into Addgene plasmid #50716). Editing efficiency was quantified by flow cytometry on a BD LSRFortessa™ instrument. The most efficient *CRX*-targeting sgRNA (AGACGTCTATGCCCGTGAGGAGG) was cloned into pLCRISPR-EFS-GFP (Addgene #57818). For shRNA experiments, *MYC*-targeting shRNA (sc-29226-SH) and scrambled shRNA control (sc-108060) were purchased from Santa Cruz.

Lentiviral particle production HEK293T cells were seeded in 100 mm dishes at a density of 1 × 10^6^ cells/cm^2^ and transfected 24h later with 6 µg of targeting plasmid (pCDH-derived, pLCRISPR-derived and shRNA-derived), 3 µg of psPAX2 (Addgene #11260) and 3 µg of pMD2.G (Addgene #12259) using FuGENE® HD Transfection Reagent (Promega) at a ratio of 2.5 µL FuGENE per 1 µg DNA in 125 µL Opti-MEM (Thermo Fisher). After 24h, the culture medium was replaced, and viral supernatants were collected every 12h over a 48h period. Collected supernatants were clarified by centrifugation and filtered through a 0.45 µm polyethersulfone (PES) filter (Millipore), then concentrated using Lenti-X™ Concentrator (Takara) at a 1:3 volume ratio overnight at 4°C. Viral pellets were recovered by centrifugation at 1,500 × g for 45 min at 4 °C, resuspended in PBS, aliquoted, and stored at -80°C until use. pCDH-derived and pLCRISPR-derived lentiviral preparations were titrated on HEK293T cells, and transduction efficiency was quantified by flow cytometry using a BD LSRFortessa™ instrument. shRNA-expressing lentiviruses were titrated on HEK293T cells following puromycin selection, and cell viability was assessed using the CellTiter-Glo® Luminescent Cell Viability Assay (Promega).

Lentiviral transduction of cell lines HDMB03 and MB3W1 cells were transduced with lentiviral particles at a multiplicity of infection (MOI) of 1 for 48h, followed by complete medium replacement. Five days post-transduction, cells were sorted for mCherry and/or GFP expression using fluorescence-activated cell sorting (100 μm nozzle). Sorted populations were expanded in culture and used for subsequent experiments. For *cMYC* shRNA transductions, an MOI of 0.5 was applied, as higher MOI of 1 resulted in reduced cell viability.

### CellTiter-Glo

Transduced HDMB03 cells were seeded in triplicate at a density of 4 × 10^6^ cells/cm^2^ in 96-well plates and incubated for 7 days at 37°C in a humidified atmosphere with 5% CO_2_. Fresh medium was added on day 4. On day 7, 100 µL of CellTiter-Glo® reagent (Promega) was added to each well and incubated for 10-15 min at room temperature, protected from light. Luminescence was measured in white plates using GloMax® Explorer Microplate Reader (Promega). A total of five independent biological replicates were performed.

Statistical analyses were performed using one-way ANOVA, followed by Dunnett’s post hoc test to compare each condition with the unmodified control cell line.

### *In vivo* tumor growth

Orthotopic xenografts were performed in 6 to 8-week-old female NOD scid gamma (NSG/NXG) mice. All animal experiments for this study were conducted according to the animal welfare regulations approved by the responsible authorities in Baden-Württemberg, Germany (Regierungspraesidium Karlsruhe, approval number: G-75/20).

HDMB03 and MB3W1 cells were harvested at 70– 90% confluence, dissociated into single-cell suspensions, and resuspended 1:1 in Corning® Matrigel® Basement Membrane Matrix (Corning) at a final concentration of 62,500 cells/µL (HDMB03) or 50,000 cells/µL (MB3W1). Suspensions were kept on ice until transplantation.

Mice were anesthetized with inhaled isoflurane (1.5% O_2_, 2% isoflurane). The surgical site was prepared by applying Bepanthen to the scalp, and Bepanthen ointment was also applied to the eyes to prevent dehydration. An approximate 1-cm incision was made along the midline from between the ears to the occipital bone, and the skull was exposed and cleaned using cotton swab. Using a stereotactic frame, a burr hole was created with a 21-gauge needle at the following coordinates relative to lambda: –1.5 mm mediolateral, –2.0 mm anteroposterior, and –2.0 mm dorsoventral. A 4 µL cell suspension was injected into the cerebellum with a 10 µL Hamilton syringe, and the needle was withdrawn slowly to prevent reflux. The incision was closed with veterinary-grade tissue adhesive (Vetbond, 3M). Following surgery, mice were recovered from anesthesia and transferred to clean cages.

Animals were monitored daily by trained staff for signs of neurological impairment or tumor-related distress. Mice developing signs were euthanized immediately by CO_2_ inhalation, and brains were collected for cryo-sectioning or paraffin embedding. Additional tumor tissue was harvested and snap-frozen for molecular analyses.

### Bulk experimental RNA-seq data acquisition

HDMB03 or MB3W1 cell pellets, as well as flash-frozen transplanted tumor tissue, were processed for RNA extraction using the RNeasy® Micro Kit (Qiagen) according to the manufacturer’s instructions. RNA integrity was assessed using the RNA ScreenTape assay on the Agilent TapeStation system, following the manufacturer’s protocol. Only samples with a RNA Integrity Number (RIN) ≥ 9.0 were retained for downstream processing. Two micrograms of total RNA per sample were submitted to the Genomics Core Facility at the German Cancer Research Center (DKFZ) for bulk RNA sequencing.

Sequencing libraries were prepared using the Illumina TruSeq mRNA stranded Kit following the manufacturer’s instructions. Briefly, mRNA was purified from 500ng of total RNA using oligo(dT) beads. Then poly(A)+ RNA was fragmented to 150 bp and converted to cDNA. The cDNA fragments were then end-repaired, adenylated on the 3′ end, adapter ligated and amplified with 15 cycles of PCR. The final libraries were validated using Qubit (Invitrogen) and Tapestation (Agilent Technologies). 2x 150 bp paired-end sequencing was performed on the Illumina NovaSeq X according to the manufacturer’s protocol. On an average 25 million paired-end reads per sample were generated.

### Protein extraction, quantification, and immunoblotting

Snap-frozen cell pellets were kept on ice and homogenized in RIPA buffer (Sigma Aldrich) supplemented with Complete™ protease inhibitor cocktail (Roche). Tissue was lysed by pipetting up and down and incubated on ice for 30 min with vortexing every 10 min. Lysates were clarified by centrifugation at 17,000 × g for 15 min at 4 °C, and supernatants were collected. Protein concentration was determined using Pierce™ BCA Protein Assay Kit (Thermo Fisher) with BSA standards (25 to 2.000 ug/ uL). Absorbance at 562 nm was measured using GloMax® Explorer Microplate Reader (Promega) and sample concentrations were obtained from calculation out of the standard curve.

For SDS-PAGE, lysates were mixed with 4× NuPAGE™ LDS Sample Buffer (Thermo Fisher) and NuPAGE™ Sample Reducing Agent (Thermo Fisher), heated at 95°C for 5 min, and centrifuged briefly. Equal amounts of protein (20–30 µg per lane) were loaded onto NuPAGE™ 4–12% Bis-Tris WedgeWell™ gels and run in Bolt™ MES SDS Running Buffer (Thermo Fisher) with NuPAGE™ Antioxidant. Electrophoresis was performed at 100V for 120 min. Proteins were transferred to PVDF membranes using the iBlot2™ system (Thermo Fisher) according to the manufacturer’s protocol. Membranes were blocked in 5% (w/v) skim milk in TBS-T for 1 h at room temperature and incubated with primary antibodies overnight at 4 °C, followed by HRP-conjugated secondary antibodies for 1 h at room temperature. Signals were detected by chemiluminescence (ECL substrate, Thermo Fisher). The following antibodies were used in this study: TRPC3 (Cell Signaling, 77934S, 1:1,000); NR2E3 (R&D, PP-H7223-00, 1:1000); EOMES (Invitrogen, 14-4877-82, 1:1,000); PAX6 (Santa Cruz Biotechnology, sc-81649, 1:500); cMYC (Cell Signaling, 9402S, 1:1,000); CRX (Santa Cruz Biotechnology, sc-377138, 1:500); b-ACTIN (Abcam, ab49900, 1:10.000).

### Immunofluorescence

Tumor-bearing brain tissue was dissected and immediately immersed in 4% paraformaldehyde at 4°C for 24 hours for fixation. Following fixation, samples were sequentially cryoprotected by immersion in 30% sucrose in PBS at 4°C, until tissues sank to prevent ice-crystal formation during subsequent freezing. Samples were embedded in optimal cutting temperature (OCT) compound in labeled cryomolds. Embedding was followed by rapid freezing of the cryomolds using dry ice until samples were completely frozen. Frozen blocks stored at -80°C until sectioning. Cryo-sectioning was performed using a cryostat set between -20°C and -24°C and 10-μm thick sections were cut and transferred onto glass slides. Slides were stored at -80°C until use.

For immunofluorescence staining of CRX and EOMES, slides were equilibrated to room temperature and washed in PBS. Brain sections were submitted to antigen retrieval by transferring slides to pre-heated citrate buffer and steamed for 20 min, followed by cooling at room temperature for 30 min. Sections were circled with a hydrophobic pen (Dako) and blocked in 10% normal donkey serum (NDS) in PBS-T for 1h. CRX (#ab140603, 1:500) antibody was prepared in blocking buffer and applied overnight at 4°C. Control slide received only blocking buffer.

Slides were washed three times in PBS-T for 10 minutes. MB3W1 sections were incubated with Anti-Rabbit AlexaFluor 488 (Life Technologies, A-31573, 1:400) and DAPI (1:1000). FlexAble 2.0 CoraLite® Plus 647 Antibody Labeling Kit for Rabbit IgG (Proteintech, KFA501) was used to label EOMES antibody and was prepared following manufacturer’s recommendations. Briefly, 6 uL of EOMES antibody (#66325) was incubated with 6 uL of FlexLinker and 36 uL of Flex Buffer and incubated at RT protected from light for 5 min. 12 uL of Flex Quencher was added and incubated for 5 min at RT protected from light. EOMES conjugated Antibody was diluted 1:250 in 10% NDS in PBS-T and incubated with CRX-stained sections overnight at 4°C. Control slides were FlexAble mock only. Images were acquired using Confocal LSM 980 AiryScan 2 (Zeiss).

